# Development and applications of chimera platforms for tyrosine phosphorylation

**DOI:** 10.1101/2023.03.05.531183

**Authors:** Rajaiah Pergu, Veronika M. Shoba, Santosh K. Chaudhary, Dhanushka N. P. Munkanatta Godage, Arghya Deb, Santanu Singha, Uttam Dhawa, Viktoriya Anokhina, Sameek Singh, Sachini U. Siriwardena, Amit Choudhary

## Abstract

Chimeric small molecules that induce post-translational modification (PTM) on a target protein by bringing it in proximity to a PTM-inducing enzyme are furnishing novel modalities to perturb protein function. Despite recent advances, such molecules are unavailable for a critical PTM, tyrosine phosphorylation. Furthermore, the contemporary design paradigm of chimeric molecules, formed by joining a non-inhibitory binder of the PTM-inducing enzyme with the binder of the target protein, prohibits the recruitment of most PTM-inducing enzymes as their non-inhibitory binders are unavailable. Here, we report two platforms to generate phosphorylation-inducing chimeric small molecules (PHICS) for tyrosine phosphorylation. We generate PHICS from both non-inhibitory binders (scantily available, platform 1) and kinase inhibitors (abundantly available, platform 2) using cysteine-based group transfer chemistry. PHICS triggered phosphorylation on tyrosine residues in diverse sequence contexts and target proteins (e.g., membrane-associated, cytosolic) and displayed multiple bioactivities, including initiation of a growth receptor signaling cascade and death of drug-resistant cancer cells. These studies provide an approach to induce biologically relevant PTM and lay the foundation for pharmacologic PTM editing (i.e., induction or removal) on target proteins using abundantly available inhibitors of PTM-inducing or erasing enzymes.

**TOC:** 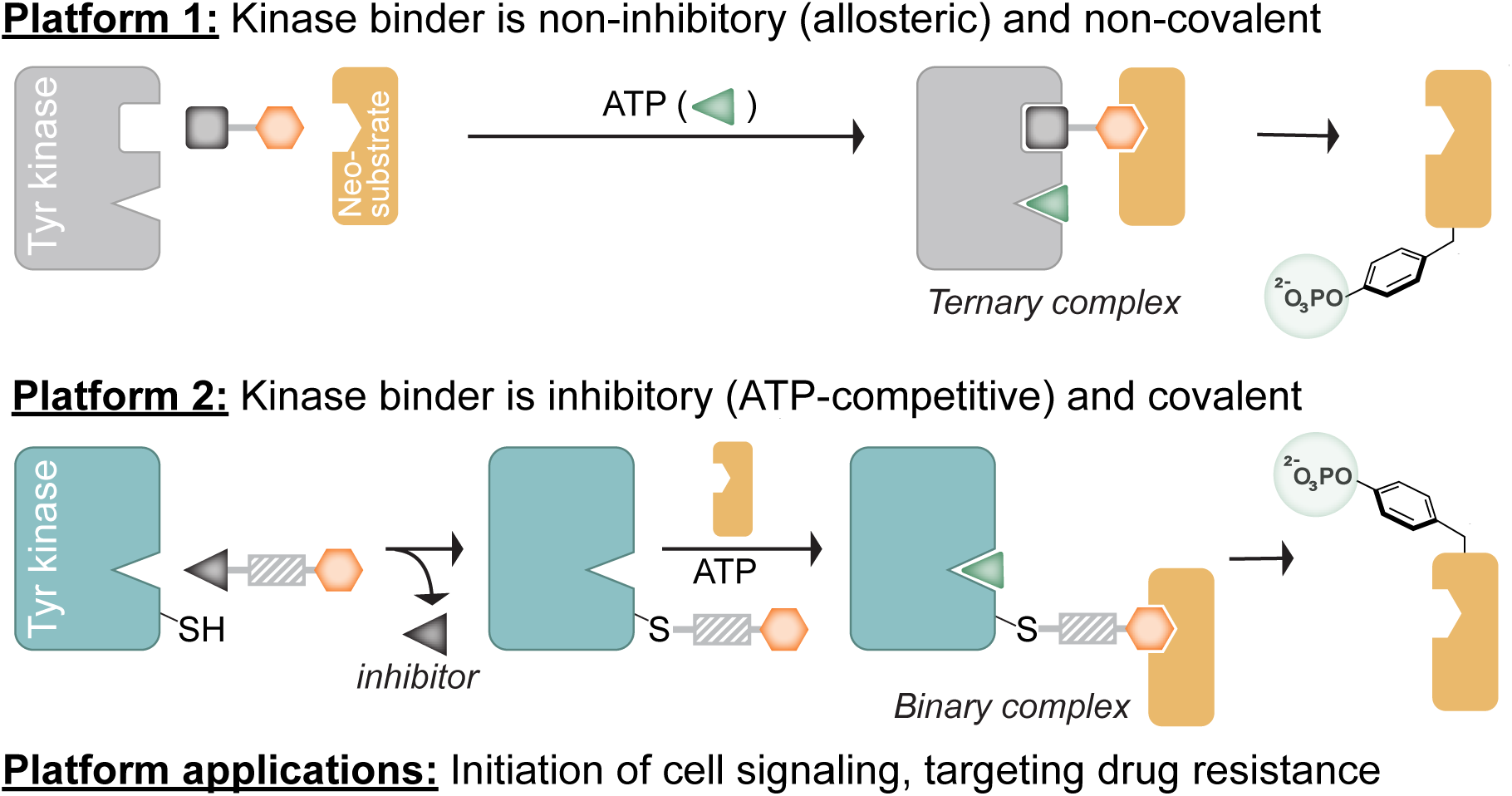

## INTRODUCTION

Protein phosphorylation is the most frequent and well-studied post-translational modification (PTMs)^1,2,3^, and its dysregulation is associated with various pathologies.^4,5^ In particular, phosphorylated tyrosine (pTyr) is vital^6,7^ in many diverse cellular functions and signaling, including highly dynamic processes often controlled using small molecules. The human genome codes 90 tyrosine kinases,^6^ 107 tyrosine phosphatases,^8^ and over hundreds of pTyr-recognition domains (e.g., SH2 domain)^9-11^ and the immune system can produce antibodies that selectively recognize pTyr over phosphorylated serine or threonine. These attributes have led to suggestions to consider pTyr a functionally separate PTM class.^12^ While molecules that inhibit tyrosine phosphorylation exist and constitute one of the largest classes of drugs^13-15^, there are no small molecules that can induce tyrosine phosphorylation on a given target of interest, even though these compounds will be of immense utility in both basic research and biomedicine.^16^

Recently, we reported on an approach to induce phosphorylation using heterobifunctional small molecules termed phosphorylation-inducing chimeric small molecules (PHICS) that recruit endogenous Ser/Thr kinases to a target protein of interest.^17-19^ These PHICS were formed by joining small-molecule ligands of targeted protein and a kinase *via* a linker. Our previously reported PHICS brought Ser/Thr kinases (AMPK or PKC) in proximity to a target protein, inducing the phosphorylation of the latter even for non-substrates (i.e., neo-substrates) of the kinase. However, these PHICS did not recruit endogenous tyrosine kinase or induce a downstream signaling event. Furthermore, these PHICS require non-inhibitory kinase binders that are unavailable for most kinases, limiting recruitment of only two of ≈538 human kinases.^20^ Beyond phosphorylation, heterobifunctional molecules that can induce or remove other PTMs, including acetylation and glycosylation, by recruiting PTM inducing or erasing enzymes are needed.^19, 21^ However, the necessity of non-inhibitory ligands for these enzymes has been a major roadblock in the development of such heterobifunctional molecules. Finally, the previously reported PHICS did not explore the covalent binders for the target protein or kinase which will have relatively simple two-body equilibrium vs. the complex three-body equilibrium displayed by non-covalent binders.^22-26^

Herein, we report tyrosine PHICS that recruit Abelson Kinase (ABL) or Bruton’s Tyrosine Kinase (BTK) to induce the phosphorylation of neo-substrates in cells. We report five non-inhibitory molecular scaffolds to recruit ABL to a target protein, and PHICS generated from these scaffolds display hallmarks of contemporarly heterobifunctional molecules, including hook effect, turnover, and ternary complex formation in cells. Importantly, we showed that ABL PHICS was able to initiate a signaling pathway by inducing phosphorylation of epidermal growth factor receptor (EGFR). Furthermore, we report a new design paradigm of a heterobifunctional molecule consisting of a BTK inhibitor connected to the target binder through a linker that covalently labels BTK with the target-protein binder while releasing the inhibitor from BTK and enabling phosphorylation of the target by BTK. We demonstrated this group transfer chemistry with two commonly employed covalent inhibitor scaffolds (i.e., aliphatic amine and aryl amine). Using BTK or ABL PHICS, we phosphorylated tyrosine residues in a diverse sequence context, including those surrounded by neutral, acidic, or basic residues. Finally, we generated ABL-BTK PHICS using a covalent inhibitor of BTK (i.e., ibrutinib) to induce its ABL-mediated phosphorylation; these PHICS effectively kill ibrutinib-resistant cancer cell lines and inhibit downstream survival signaling of BTK, highlighting the therapeutic potential of this class of molecules.

## RESULTS AND DISCUSSION

### Identification of ABL binders for generation of BRD4 PHICS

For the tyrosine kinase recruited by PHICS, we first chose ABL, as it can phosphorylate wide-ranging target sequences and has several non-inhibitory, selective, and allosteric binders that also have co-crystal structures to aid the rational design of PHICS.^27-30^ We selected four scaffolds that competitively bind to the myristoyl pocket of ABL: dihydropyrazole (scaffold 1, Figure S1A), thiazole (scaffold 2, Figure S1B), pyrazole (scaffold 3, Figure S1C),^30^ and hydantoin (scaffold 4, Figure S2).^29^ Using molecular docking studies, we identified the solvent-exposed 3-amino group as the linker attachment site for the dihydropyrazole, pyrazole, and thiazole scaffolds (Figure S1A‒C, PDB ID: 6NPV and 6NPE).^30^ For the hydantoin scaffold (PDB ID: 3PYY),^29^ linkers were grown from the *ortho, meta*, or *para* positions of the solvent-exposed aryl ring (Figure S2). Previous reports for synthesizing hydantoin scaffold^29^ involving the Vilsmer-Haack reaction provided poor yield^31^ and were sensitive to substitutions on the aryl ring (Figure S3A) preventing exploration of linker growth from -*ortho*, -*meta*, or -*para* positions from aryl ring. We developed a new synthetic route with fewer steps to rapidly access hydantoin scaffold precursors by involving Cu-mediated high-yielding C─N bond coupling^32^ between pyrazole and appropriately substituted phenylboronic acids (Figure S3B). By applying this synthetic strategy, we were able to rapidly generate a library of hydantoin-based PHICS with linkers through the -*ortho*, -*meta*, and -*para* positions of the aryl ring.

To assess the efficacy of these scaffolds, we synthesized four different PHICS, each containing one of the four binders of ABL, a linker comprised of four polyethylene glycol units, and the small molecule (*S*)-JQ1 (Figures S3C and S4), which binds our proof-of-concept target bromodomain-containing protein 4 (BRD4).^33^ With these compounds in hand, we measured PHICS-induced phosphorylation using recombinant ABL and BRD4 proteins. Dihydropyrazole induced the most BRD4 phosphorylation followed by PHICS based on thiazole, while the pyrazole scaffold (an oxidized form of the dihydropyrazole) induced significantly less phosphorylation (Figure S5). Hydantoin-derived PHICS induced less phosphorylation than the dihydropyrazole-derived PHICS with the meta-substituted analog showing the highest phosphorylation levels compared to the ortho- and para-substituted analogs (Figures S5 and S6). These studies identified the dihydropyrazole scaffold as the most efficient for inducing BRD4 phosphorylation.

To more deeply characterize the ability of the dihydropyrazole scaffold to induce BRD4 phosphorylation, PHICS **1** (Figure 1A) was selected for further studies together with three controls: **2** generated by replacing (*S*)-JQ1 in **1** with inactive enantiomer (*R*)-JQ1, **3** formed by replacing the dihydropyrazole in **1** with pyrazole scaffold, and **4** a dihydropyrazole scaffold with a short linker. In biochemical settings, we confirmed that tyrosine phosphorylation of BRD4 occurred only in the presence of all components of the ternary complex, namely **1**, BRD4, and ABL (Figure 1B). We then assessed the formation of the ternary complex between **1**, BRD4-GST, and ABL-His was assessed using Amplified Luminescent Proximity Homogenous assay (AlphaScreen),^34^ wherein **1**, but not **2**, showed a “hook-effect” which is a hallmark of 3-body equilibrium^22, 23^ (Figure 1C). Here, as the concentration of **1** increases, there is an increase in the population of the ternary complex. However, after a certain concentration, the binary complexes between **1** and ABL and **1** and BRD4 dominate the equilibrium over the ternary complex, resulting in a hook-shape dose curve.

**Figure 1.**
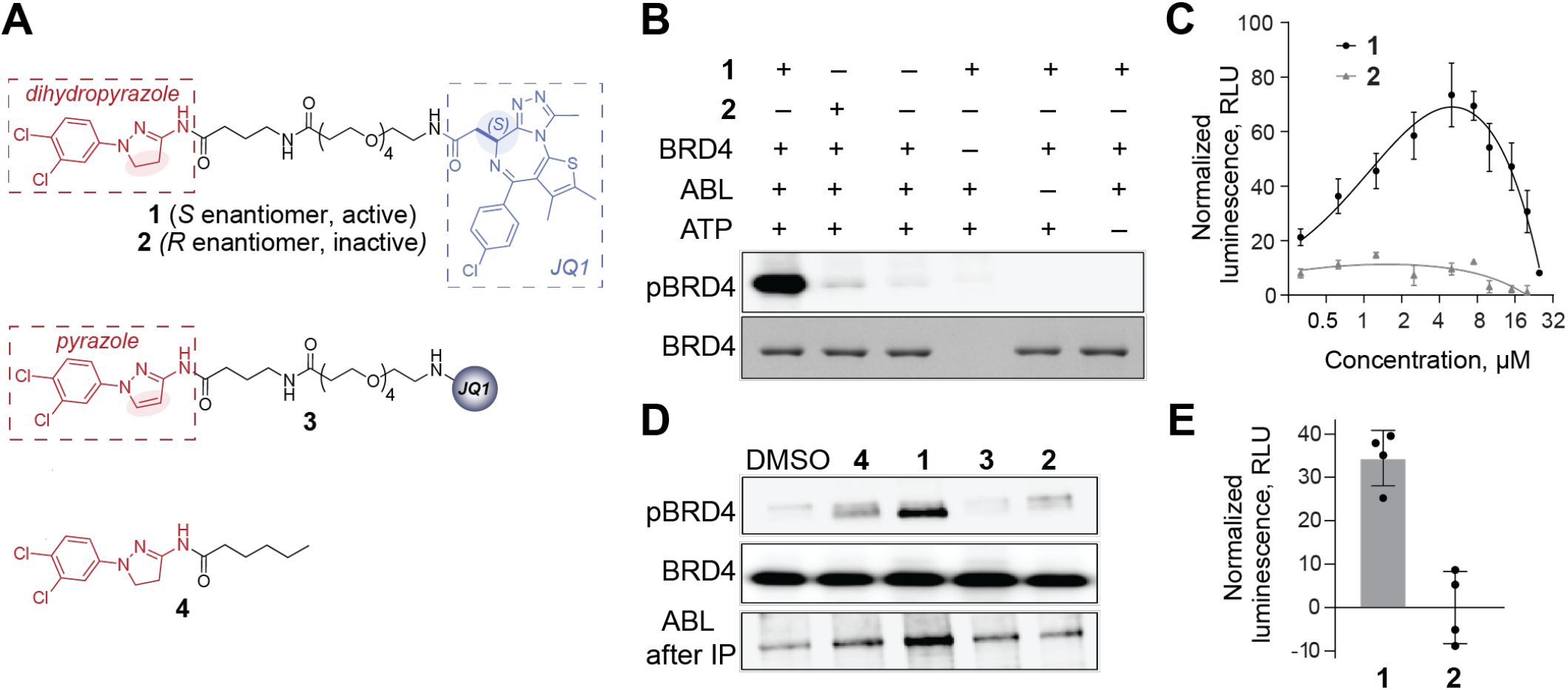
**(A)** Structures of ABL-BRD4 PHICS **1** and control compounds **2-4. (B)** PHICS-induced phosphorylation of BRD4 by ABL *in vitro*. **(C)** PHICS-induced ternary-complex formation between BRD4 and ABL observed by Alpha Screen assay. **(D)** PHICS-induced phosphorylation of BRD4 by ABL in HEK293T cells; co-immunoprecipitation (Co-IP) of ABL with BRD4 in the presence of PHICS. **(E)** ADPGlo assay for BRD4 phosphorylation by ABL in the presence of **1** and its inactive isomer **2**.

To demonstrate BRD4 phosphorylation in cells using PHICS, we used a construct of *BRD4* lacking an intrinsic nuclear localization signal^35^ and incubated HEK293T cells transfected with both *HA-BRD4* and *ABL-FLAG* plasmids with bifunctional molecules and controls. Here, HA-based immunoprecipitation (IP) showed significantly higher co-immunoprecipitation of ABL-FLAG with **1** compared to the control compounds (i.e., **2, 3**, or **4**), suggesting ternary-complex formation is specific to **1** (Figure 1D). Probing the immunoprecipitated HA-BRD4 with a pan phospho-tyrosine antibody showed significantly higher phosphorylation levels in cells treated with **1** than in the control compounds, confirming that the observed phosphorylation is arising from PHICS-mediated ternary complex formation between ABL and BRD4. To confirm that the reversible binding of PHICS to both BRD4 and ABL allows it to exhibit catalytic turnover, we used an ADP-Glo assay^36^ to determine the amount of ADP generated per molecule of ABL using active molecule **1** and control **2** (Figure 1E). Here, the amount of ADP generated in the reaction with **1** was 2261 ± 411 nM higher than the amount of ADP generated in the reaction with **2** at a limiting concentration of ABL kinase (30 nM), confirming that PHICS exhibits turnover, like our previously reported Ser/Thr PHICS.^17^

### PHICS can activate EGFR signaling

EGFR is a tyrosine kinase receptor that responds to the epidermal growth factor (EGF) binding to initiate a signaling pathway that regulates the growth, proliferation, survival, and differentiation of cells.^37, 38^ Upon EGF binding, EGFR oligomerizes and autophosphorylates tyrosine residues, which acts as docking site of proteins with phosphotyrosine-binding SH2 or PTB domains leading to a series of molecular events and cellular response.^39^ We were interested in determining if ABL PHICS can trigger signaling by phosphorylating relevant EGFR tyrosines. Some kinases prefer specific residues surrounding the phosphosite while others do not. Since signaling relevant EGFR tyrosines are present in diverse sequence environments (i.e. surrounded by acidic, basic, or neutral residues), we were interested in determining if ABL PHICS can phosphorylate these diverse tyrosines and trigger signaling. We constructed a catalytically inactive intracellular domain of the EGFR variant fused to FKBP^F36V^ (*iEGFR-FKBP*^*F36V*^*-FLAG*), and synthesized PHICS **5** using the FKBP^F36V^ binder AP1867 (Figure 2A, S7).^40^ HEK293T cells transfected with *iEGFR-FKBP*^*F36V*^*-FLAG* and *ABL-HA* were treated with **5** or DMSO. Immunoblotting of cell lysates and probing with site-specific phospho-tyrosine antibodies suggested that ABL PHICS could phosphorylate tyrosines present in diverse sequence environments (Figure 2B) in line with previous reports that ABL lacks sequence motifs.^28, 41^ Similarly, we observed phosphorylation when using catalytically inactive HER2, another receptor tyrosine kinase (Figure S8), pointing to the generality of the designed PHICS.

**Figure 2.**
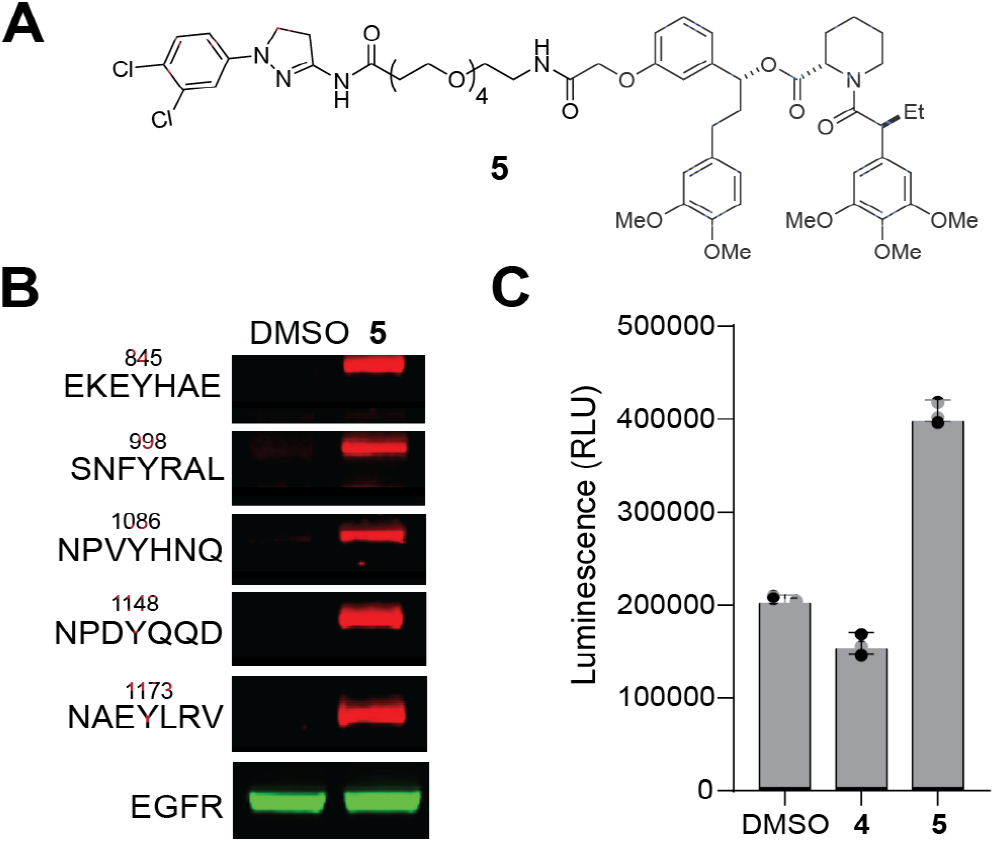
**(A)** Structure of ABL-FKBP^F36V^ bifunctional molecule **5** used in PHICS-induced phosphorylation of EGFR-FKBP^F36V^ construct. **(B)** Diverse sequence environments of Tyr phosphorylated by ABL in cells in the presence of **5** detected using antibodies specific for phospho-EGFR. **(C)** Induction of luciferase-based reporter gene with serum response element by PHICS **5**, but not by DMSO or ABL binder **4**.

To evaluate the effect of PHICS **5** on downstream EGFR signaling, we used a previously described luciferase-based reporter gene assay with serum response element (SRE).^42^ Here, HEK293T cells were co-transfected with *iEGFR-FKBP*^*F36V*^*-FLAG* and pGL4.33[luc2P/SRE/Hygro] vectors, treated with PHICS **5**, or controls (DMSO or **4**); the activation of EGFR signaling was quantified using Promega’s ONE-Glo™ Luciferase Assay System. PHICS **5** induced a two-fold higher luciferase activity than DMSO or ABL binder **4**, indicating activation of EGFR by **5** (Figure 2C). These studies confirm that cytoplasmic ABL can be recruited to membrane-localized targets to induce functionally relevant tyrosine phosphorylations flanked by various amino-acid sequences and further confirms the broad range of phosphorylating motifs available to ABL kinase.^43, 44^

### PHICS can be developed from covalent binder of the target protein

While covalency at the kinase end allows phosphorylation of multiple molecules of the target protein per molecule of PHICS owing to turnover, the use of covalent binder of the target protein will block such turnover by PHICS and may dramatically reduce the efficacy of PHICS to induce phosphorylation. We were interested in determining if efficacious PHICS can be generated using a covalent binder of the target protein. For this purpose, several bifunctional molecules were designed by connecting hydantoin or dihydropyrazole binders of ABL to chloroalkane linkers that covalently bind HaloTag (Figure 3A and S9).^45^ To confirm the cell permeability of these compounds and their ability to efficiently label HaloTag, we used a competition assay wherein HEK293T cells stably expressing HaloTag were treated with compounds **7, 9, 11**, or **12** and controls (**10** and **4**), and the lysates were subsequently labeled with tetramethylrhodamine (TMR)-HaloTag, a fluorescent probe.^46^ We found that the lysates from cells pretreated with HaloTag-PHICS were not labeled by TMR-HaloTag, whereas control-treated lysates (**10** and **4**) showed efficient TMR-labelling (Figure S10), indicating that HaloTag-PHICS are cell-permeable and efficiently label the HaloTag protein. To assess the relative ability of these covalent PHICS to induce tyrosine phosphorylation, we transiently expressed the ABL-HA construct in HEK293T cells stably expressing the HaloTag protein and treated these cells with various covalent PHICS (compounds **7, 9, 11**, and **12**) and controls (compounds **6, 8, 10**, and **4**). We observed significant tyrosine phosphorylation on HaloTag in the presence of active PHICS compared to control-treated samples (Figure 3B). These results demonstrate that irreversible binding is tolerated by both the kinase and the target to generate a functional PHICS, opening avenues for the use of covalent inhibitors, a burgeoning class of chemical matter.^47^

**Figure 3.**
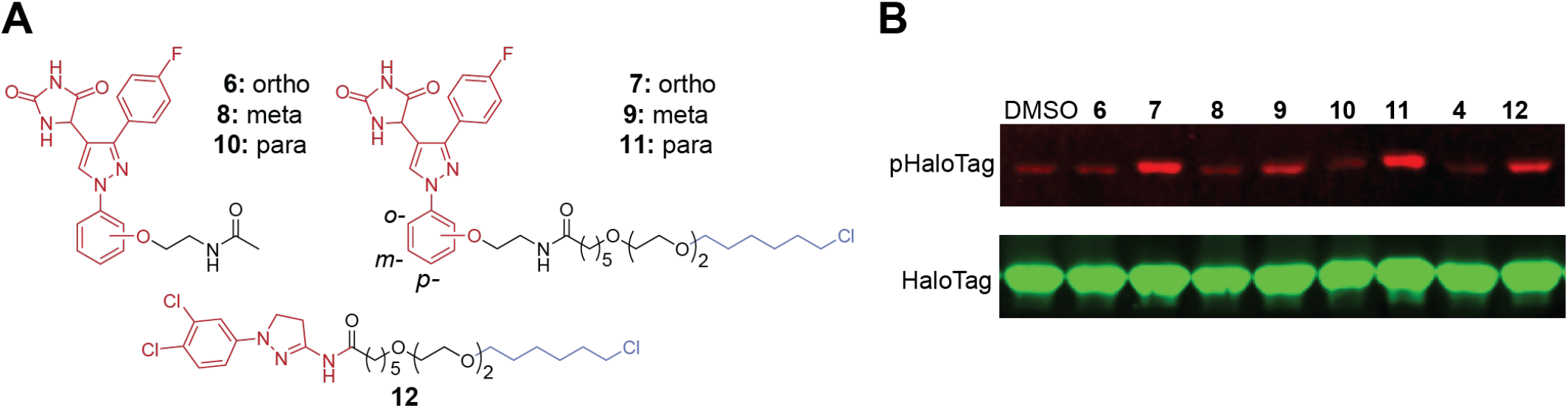
**(A)** Structures of hydantoin and dihydropyrazole-derived HaloTag bifunctional and control compounds. **(B)** PHICS-induced phosphorylation of HaloTag by ABL in cells.

### PHICS can be developed using kinase inhibitors

All the reported PHICS were developed using non-inhibitory allosteric binders, which are rare, though kinase inhibitors are plentiful and form an important class of therapeutic.^48^ A PHICS can be designed^49-51^ from a kinase inhibitor in which a nucleophile (e.g., cysteine) near the inhibitor-binding pocket can initiate a group transfer chemistry^52-59^ resulting in appending the target-protein binder to the kinase while releasing the inhibitor from the kinase’s active site (Figure 4A). The kinase binder in such cysteine-triggered PHICS can be derived from abundantly available acrylamide-based inhibitors, where the amide nitrogen is often an aryl-amine^60^ (**13**, Figure 4B) or less frequently aliphatic amine^61^ (**14**, Figure 4B).^47^ Using these BTK inhibitor scaffolds, we synthesized BRD4 PHICS **17** (aryl amine scaffold, Figure 4B) and **19** (aliphatic amine scaffold, Figure 4B), wherein the inhibitor is connected to the BRD4 binder JQ1 via a cleavable methacrylamide linker (Figure S11). We also synthesized inactive controls (**18** and **20**, Figure 4B) for BRD4 using (*R*)-JQ1 and tested whether **17** or **19** can rewire BTK’s specificity and induce the phosphorylation of BRD4. Indeed, we observed much higher BRD4 phosphorylation in the presence of **17** (Figure 4C) or **19** (Figure 4D) than with inactive controls (**18** or **20**) in cells. Using the same assessments of ternary-complex formation as previously, we observed significantly higher levels of co-immunoprecipitated BTK-FLAG and higher levels of BRD4 phosphorylation in samples treated with **17** or **19** compared to samples treated with inactive controls, **18** or **20** (Figures 4C-D).

**Figure 4.**
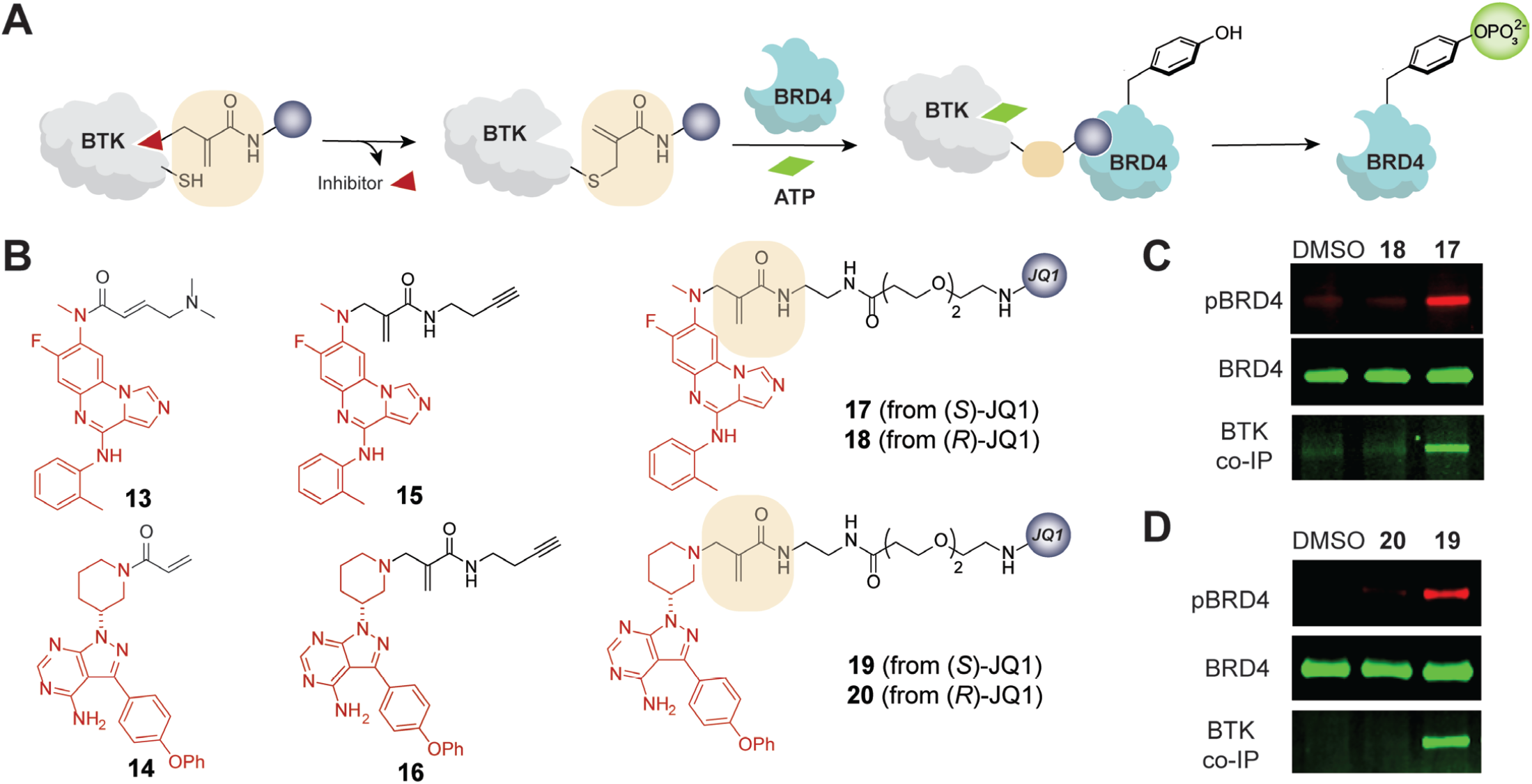
**(A)** Schematic representation of BTK tagging via inhibitor-directed addition-elimination, ternary complex formation, and phosphorylation of BRD4. **(B)** Structures of known BTK inhibitors **13** and **14**, corresponding alkyne-containing analogs **15** and **16**, and BTK-BRD4 PHICS **17** and **19** with their inactive analogs **18** and **20. (C-D)** PHICS-induced phosphorylation of BRD4 by BTK in cells and co-IP of BTK-FLAG with HA-BRD4 upon treatment with **17**-**18** (**C**) or **19**-**20** (**D**).

Next, we performed target engagement and labeling studies designed to support the mechanism outlined in Figure 4A. Using compounds that have an alkyne handle in place of JQ1 (**15** and **16**, Figure 4B), we confirmed that the scaffolds of **17** or **19** (Figure 4B) covalently engaged BTK in cells. Briefly, HEK293T cells transiently expressing BTK were treated with alkyne-containing compounds **15** and **16** for 4 h. After washing with PBS, Cu-catalyzed click reaction was performed on the lysates using sulfo-Cy5.5 azide; BTK labeling was confirmed via in-gel fluorescence (Figure 5A). To demonstrate that the BTK inhibitor scaffold is released from the ATP-binding pocket, we used a reporter assay based on Bioluminescence Resonance Energy Transfer (BRET)^62^ between a nanoluciferase (nanoLuc) and a fluorophore probe that binds to the ATP pocket; a higher inhibitor occupancy in this ATP pocket will prevent the binding of the tracer, and lower the BRET signal^63^ (Figure 5B). As expected, **13** and **14** dramatically lowered the BRET signal indicating blockage of the ATP pocket, while their derivatives with cleavable methacrylamide linkers (alkynes **15** and **16**, BRD4-BTK PHICS **17** and **19**) did not affect the binding of tracer (Figure 5C-D). Agreeing with these findings, we observed higher autophosphorylation of BTK with PHICS compounds **17** and **19** compared to the parent inhibitors **13** and **14** (Figure S12). These studies demonstrate that PHICS can be generated from cysteine-targeting covalent inhibitors, which are available for many kinases^47, 64^ with diverse cellular and tissue localization, sequence preferences, and successful clinical outcomes.

**Figure 5.**
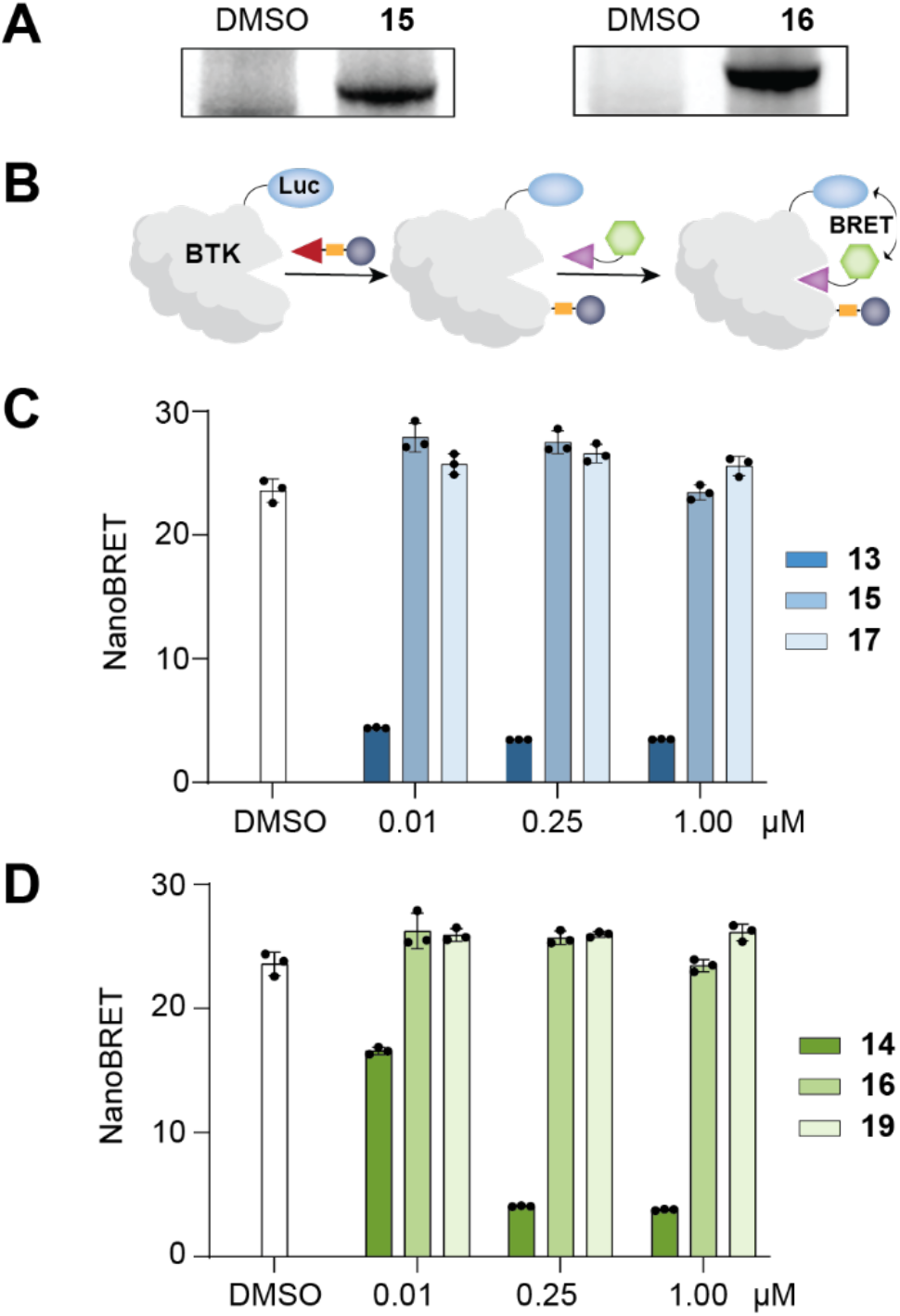
**(A)** Demonstration of covalent labeling of BTK in cells by **13** and **14** at 1 µM via in-gel fluorescence. **(B)** Schematic representation of nanoBRET assay for evaluation of BTK binding pocket occupancy. **(C-D)** NanoBRET assay for binding pocket occupancy for known BTK inhibitors **13** (**C**) and **14** (**D**), and their derivatives designed to label BTK via group transfer chemistry.

### PHICS induce death of drug-resistant cancer cells

After validating two classes of Tyr-PHICS (ABL- and BTK-derived), we explored the functional capabilities of bifunctional molecules consisting of binders of both kinases since these kinases are often overexpressed in cancer cells, and we have reported that Ser/Thr PHICS can induce inhibitory phosphorylation on BTK.^18^ We designed compound **21**, connecting the reversible binder of ABL with the covalent binder of BTK via a PEG3 linker (Figure 6A and S13) and used this PHICS along with a mixture of the separate binders (**22** and **4**) for the treatment of HEK293T cells co-transfected with *ABL-HA* and *BTK-FLAG* plasmids. Western blotting with a pan anti-phospho-tyrosine antibody revealed increased levels of BTK phosphorylation in the presence of **21** when compared to the control (mixture of binders **22** and **4**) (Figure 6B). Surprisingly, the level of ABL phosphorylation remained unchanged. This phosphorylation outcome motivated us to evaluate BTK-ABL bifunctional **21** in cancer cell lines that depend on ABL or BTK, particularly those that were resistant to the known BTK-targeting drug, ibrutinib^65^. When tested in BTK-dependent ibrutinib-resistant Mino and Raji cell lines, **21** reduced their viability with an EC_50_ of 1.7 µM and 4.9 µM, respectively (Figure 6C).^65^ In contrast, **21** did not impact the viability of BCR-ABL dependent K562 cells (Figure S14), in agreement with our expectation based on the unaffected phosphorylation of ABL in HEK293T-based studies (Figure 6B). We noted that neither the mixture of individual binders (**22** and **4**) nor ibrutinib induced the same effect of cell viability in BTK-dependent and ibrutinib-resistant cancer cells as compound **21**. These results with ibrutinib are especially interesting, as this BTK inhibitor is an approved therapeutic agent for several B-cell cancers. However, the emergence of resistance has resulted in a need for next-generation therapeutics;^66^ the promising activity of BTK-ABL PHICS in ibrutinib-resistant cell lines opens such an alternative therapeutic strategy.

**Figure 6.**
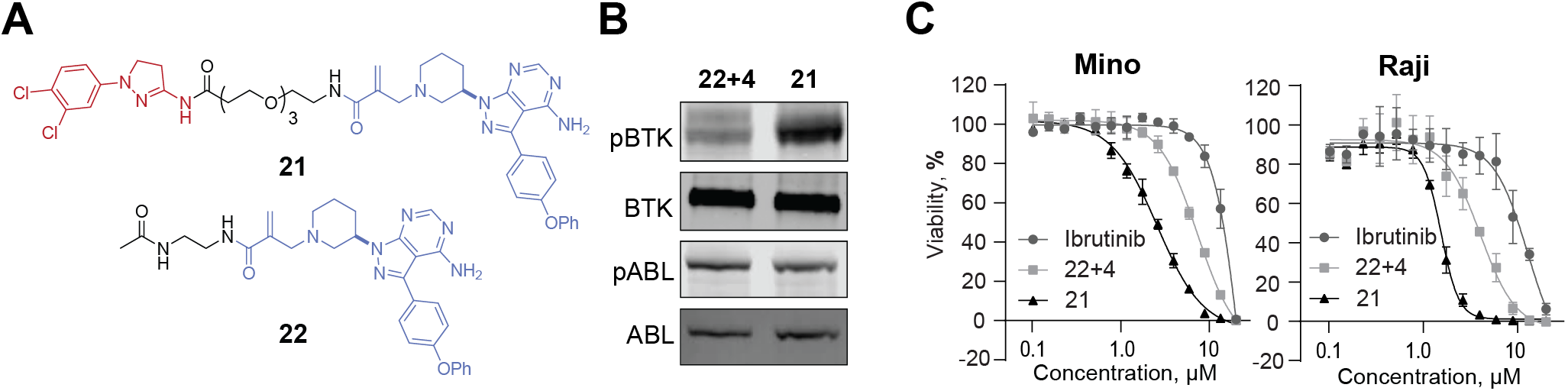
**(A)** Structures of ABL-BTK bifunctional molecule **21** and BTK binder **22. (B)** PHICS-induced phosphorylation of BTK by ABL in HEK293T cells. **(C)** Effect of PHICS **21** on the viability of Mino and Raji cells.

## CONCLUSION

To address the lack of technologies for the facile editing of PTMs (i.e., addition or removal) on a given protein of interest, we report a new class of bifunctional molecules that induce functionally relevant tyrosine phosphorylation by recruiting ABL or BTK. PHICS triggered activation of EGFR, which belongs to the receptor tyrosine kinase superfamily that includes insulin receptor. A PHICS that similarly activates insulin receptor signaling may furnish an orally available, small molecule substitute for the insulin that requires injection and may offer an alternative therapeutic modality to patients with insulin resistance. Furthermore, PHICS induced death of drug-resistant cancer cells, offering a potentially novel and alternative therapeutic modality to kinase inhibitors against which resistance has developed. The CRISPR-Cas system^67, 68^ has furnished technologies for facile editing of the DNA or RNA, but methods for facile editing of post-translational modifications (i.e., addition or removal) on a given protein of interest is still challenging. We report an approach to induce phosphorylation on the target protein by generating PHICS that utilize kinase inhibitors and cysteine-triggered group transfer chemistries. Since inhibitors for several PTM-inducing or removing enzymes are available, and group transfer chemistries can be implemented using other nucleophilic residues (e.g., lysine, tyrosine, methionine);^69-71^ these studies lay the foundation for pharmacologic editing of PTMs on proteins. Overall, these studies further highlight the power of bifunctional molecules to endow neo-functions to proteins in cells with value in basic research and medicine.

## ACKNOWLEDGMENTS

We thank Dr. Mary O’Reilly (Pattern, Broad Institute) for contributive discussions and help with figure design. This work was supported by the Merkin Institute of Transformative Technologies in Healthcare, Mark Foundation, Burroughs Wellcome Fund (Career Award at the Scientific Interface), DARPA (N66001-17-2-4055 and HR0011-21-2-0010), and NIH (R21AI154099). V.M.S. was supported by Damon Runyon Postdoctoral Fellowship.

## AUTHOR CONTRIBUTIONS

These authors contributed equally to this work.

