## Supporting Information for "Development and applications of chimera platforms for tyrosine phosphorylation"

**1. 10. Immunoblotting studies to confirm PHICS-mediated phosphorylation of Halo-Tag by ABL in HEK293T cells. 9**

### 1. Materials and methods.

#### 1. 1. Materials and general procedures.

BRD4-GST ((BD1 and BD2 (49-460)) (Cat#31044) was purchased from BPS Bioscience. ADP-Glo™ Kinase Assay kit (Cat#V6930) was purchased from Promega Corporation. OptiPlate-384 (White Opaque 384-well Microplate, Cat#6007290), Corning® 96-well plates with Solid White Flat Bottom (Corning cat#3917), Nickel Chelate AlphaLISA Acceptor Beads (Cat#AL108C), and Alpha Glutathione Donor beads (Cat#6765300) were all purchased from Perkin Elmer. Kinase assay buffer III 5x (Cat#K03-09) and Abltide peptide (EAIYAAPFAKKK, Cat#A02-58) were purchased from SignalChem. pcDNA3.1(+) HA-BRD4 (aa 49-460) and pcDNA3.1(+) ABL1b-FLAG full length plasmids (cat# OHu28559, NM_007313.2) were purchased from GenScript. BCR-ABL full length (P210 pcDNA3, plasmid # 27481) was purchased from Addgene. TransIT®-293 and TransIT®-LT1 Transfection Reagent was purchased from Mirus (MIR 2700 and MIR 2300). cOmplete™, Mini, EDTA-free Protease Inhibitor Cocktail (Cat#04693159001) and Inhibitor, Phosphatase, PhosSTOP (Cat#4906837001) were purchased from Millipore Sigma. M-PER™ Mammalian Protein Extraction Reagent (Cat#78501), Pierce™ BCA Protein Assay Kit (Cat#23225), Pierce™ Anti-HA Magnetic Beads (88837), Pierce™ Anti-DYKDDDDK Magnetic Agarose (A36797), Dynabeads™ Protein A for Immunoprecipitation (Cat# 10001D), and NuPAGE 4-12% Bis-Tris Protein Gels (Cat#NP0336 or Cat#NP0335), Novex™ WedgeWell™ 4 to 12%, Tris-Glycine (Cat#XP04120), Novex™ Tris-Glycine SDS Running Buffer (10X) (Cat#LC2675) and ABL1 protein (P3049) were purchased from Invitrogen. SuperSignal™ West Pico PLUS Chemiluminescent Substrate was purchased Thermo Scientific). Cell titer Glo reagent used for cell viability assay was purchased from Promega (Cat#G7572). NanoBRET™ Target Engagement Intracellular Kinase Assay kit, Adherent Format (cat#N2641); FuGENE HD Transfection Reagent (cat# E2311), BTK Kinase-NanocLuc fusion vector (cat#N2441) were purchased from Promega. Opti-MEM I (cat# 11058-021) was purchased from Life Technologies. pGL4.33[luc2P/SRE/Hygro] Vector was purchased from Promega (Cat#E1340). ONE-Glo™ Luciferase Assay System used for SRE reporter assay was purchased from Promega (Cat#E6110). Mino, Raji, and K562 cells were purchased from ATCC (Cat# CRL-3000, CCL-86, and CRL-3343) and grown in RPMI media supplemented with 10% FBS and 1% Penicillin-Streptomycin.

For chemical synthesis, all reagents were purchased and used as received from commercial sources without further purification. Reactions were performed in round-bottom flasks stirred with Teflon®-coated magnetic stir bars. Moisture and air-sensitive reactions were performed under a dry nitrogen/argon atmosphere. Moisture and air-sensitive liquids or solutions were transferred via nitrogen-flushed syringes. Organic solvents were degassed by bubbling nitrogen/argon through the liquid, as needed. Flash column chromatography was performed using silica gel (60 Å mesh, 20–40 μm) on a Teledyne Isco CombiFlash Rf system. HPLC purification was performed on Teledyne ISCO ACCQPrep™ HP150 equipped with an XBridge BEH Prep Column (19 mm x 250 mm). UPLC-MS was performed on a Waters ACQUITY UPLC I-Class 15 PLUS system with an ACQUITY SQ Detector 2. Nuclear magnetic resonance (NMR) spectra were collected on a Bruker AVANCE III HD 400 MHz spectrometer at room temperature (1H NMR, 400 MHz; 13C, 101 MHz). $^{1}$H and $^{13}$C chemical shifts are indicated in parts per million (ppm) and internally referenced to residual solvent signals. NMR solvents were purchased from Cambridge Isotope Laboratories, Inc., and NMR data were obtained in $CDCl_3$, $CD_3OD$, and DMSO-d6. Data for

$^{1}H$ NMR are reported as follows: chemical shift value in ppm, multiplicity (s = singlet, br s = broad singlet, d = doublet, t = triplet, dd = doublet of doublets, and m = multiplet), integration value, and coupling constant value in Hz. HRMS was performed on an Agilent 6210/6220 ESI-TOF.

#### 1. 2. Evaluation of the turnover efficiency of Tyrosine PHICS molecules by ADP-Glo assay.

ADP-Glo assay was performed in the presence of active or inactive PHICS to characterize the turnover efficiency of PHICS molecules. Kinase reaction was performed in a 96-well plate (white, flat bottom) in the presence of tested molecules, BRD4-GST, ABL1 kinase, and ATP. 10 µM of active PHICS or inactive PHICS was incubated with 30 nM ABL kinase, 1000 nM BRD4-GST (BD1 and BD2) and 100 µM ATP in the kinase assay buffer (40 mM Tris-HCl pH 7.5, 20 mM MgCl2, 0.1 mg/ml BSA, 50 µM DTT, 1% DMSO) for 2 hours at room temperature. Then, the ADP-Glo assay was performed following Promega's protocol for ABL1 kinase. Briefly, ADP-Glo reagent was introduced to the kinase reaction mixture in a 1:1 ratio and further incubated for 40 min at room temperature. After that, kinase detection reagent was added to the reaction mixture with a 1:2 ratio and maintained the reaction for another 30 min at room temperature before recording the luminescence by Envision 2104 plate reader (PerkinElmer).

The luminescence signal for BRD4 phosphorylation by active PHICS was determined by subtracting the signal from inactive PHICS. To convert the luminescence signal to the amount of ADP generated during the kinase reaction, we prepared the 25 µM ATP-to-ADP conversion curve (luminescence (RLU) versus % ATP to ADP conversion) following the Promega protocol and plotted data using GraphPad PRISM version 9.3.0. The actual ADP generation in the presence of active PHICS was calculated using the conversion curve and turnover efficiency data was plotted using GraphPad PRISM version 9.3.0 normalized to the inactive PHICS control.

#### 1. 3. Characterization of ternary complex formation (kinase-PHICS-target protein) by AlphaScreen assay.

To evaluate the ternary complex formation between ABL1 kinase, PHICS, and BRD4 target protein, we performed an AlphaScreen assay following the PerkinElmer specifications. The assay was performed in a white opaque 384-well microplate. Initially, concentration series of PHICS with DMSO control were prepared using the dilution assay buffer (50 mM HEPES pH 7.4, 150 mM NaCl, 0.1% w/v BSA, 0.01% v/v Tween 20) and maintained DMSO as 1% in final mixture. Then 7 nM ABL kinase with 6x His tag and 67 nM BRD4 (BD1 and BD2) with GST tag were introduced to the reaction mixture and incubated for 1 hour at room temperature before adding the AlphaLISA beads. Nickel Chelate AlphaLISA Acceptor beads (PerkinElmer) which can be used to capture His-tagged proteins and Alpha Glutathione Donor beads (PerkinElmer) which can be used to capture GST-tagged proteins were added to the mixture in a final concentration of 20 µg/mL and the reaction was kept at room temperature for another hour before the luminescence was recorded on Envision 2104 plate reader (PerkinElmer). The luminescence signal was further processed using DMSO control in order to obtain the normalized values. Data were analyzed and plotted using GraphPad PRISM version 9.3.0

#### 1. 4. Immunoblotting analysis to confirm the PHICS-induced tyrosine phosphorylation of BRD4 in vitro.

Proximity-mediated tyrosine phosphorylation on BRD4 was confirmed by western blotting studies using an anti-phosphotyrosine antibody followed by a kinase reaction in the presence of PHICS. Kinase reaction was performed in an assay buffer with 40mM Tris-HCl, pH 7.4, 20mM $MgCl_{2,}$ 0.1mg/ml BSA and 100μM DTT (1x kinase assay buffer III, SignalChem). PHICS, ABL kinase, BRD4 (BD1 and BD2), and ATP were diluted in the assay buffer to maintain a final concentration of 10 μM (DMSO final concentration 1%), 30 nM, 1000 nM and 100 μM respectively. Afterward, the kinase reaction mixture was incubated for 2 hours at room temperature before quenching with an SDS loading buffer. Proteins were resolved by NuPAGE 4-12% Bis-Tris protein gels and transferred to a PVDF membrane using a dry transfer iBLOT2 system. After the transfer membrane was incubated in the blocking buffer (5% milk in TBST) for 1 hour at room temperature. After that membrane was incubated with the anti-phosphotyrosine antibody, clone 4G10 primary antibody (Millipore Sigma, Cat#05-321) (1:1000) in the same blocking buffer. Following overnight incubation at 4°C, the membrane was washed 3X times with TBST buffer (TBS with 0.1% tween 20) and further incubated with HRP-conjugated secondary antibody (mouse, CReferencesell Signaling, Cat#7076) for 1 hour at room temperature. After that, the membrane was washed 5X with TBST buffer before visualizing the bands by chemiluminescence (Azure Biosystems C600 imager).

Similar method and similar concentrations of PHICS, BRD4, and ABL kinase were used to evaluate the different cores of pyrazole containing PHICS, different linkers of pyrazole containing PHICS, DPH containing PHICS with different exit vectors, and different linkers of PHICS with DPH as the ABL kinase binder.

#### 1. 5. Confirmation of ternary complex (ABL:PHICS:BRD4) formation in HEK293T cells.

Ternary complex formation (ABL kinase:PHICS:BRD4 target protein) was validated in HEK293T cells by co-immunoprecipitation studies after overexpression of ABL kinase and truncated BRD4 (aa 49-460, without nuclear localization signal) which is predominantly localized to the cytoplasm. HEK293T cells were cultured in DMEM supplemented with 10% FBS, penicillin (100 units/mL), and streptomycin (100 μg/mL). Cells were maintained at 37 ºC in a 5% CO2 humidified atmosphere.

Initially, ABL-FLAG and HA-BRD4 plasmids were transfected into HEK293T cells using TransIT®-293 transfection reagent, and cells were maintained for another 36 hours before incubation with compounds. Then cells were treated with 10 μM concentration of ABL binder, active PHICS, inactive PHICS, and DMSO control for 4 hours at 37 ºC in DMEM media with the serum. After 4 hours of incubation, cells were washed with cold PBS 1X time and lysed on ice using the cold lysis buffer (M-PER™ Mammalian Protein Extraction Reagent, EDTA-free Protease Inhibitor Cocktail (cOmplete™) and PhosSTOP). Pierce™ BCA Protein Assay Kit was used to determine the protein concentration of cell lysates, and immunoprecipitation (IP) was performed using the volume of cell lysate that contains one milligram of protein for each IP sample.

For the co-immunoprecipitation studies, HA-tagged BRD4 was immobilized on beads, and elution was probed for the FLAG-tagged ABL1b. Anti-HA magnetic beads were used, and the immunoprecipitation protocol from Thermo Scientific Pierce was followed. Briefly, 25 µL of beads were used for each sample, and beads were washed with M-PER™ buffer 2X times before incubation with cell lysates. A volume of cell lysate containing approximately one milligram of proteins was incubated with the beads at 4 ˚C overnight on a rotating shaker. The next day, beads were washed 3X times with TBST (0.05% tween 20) buffer and one time with water before elution of proteins. SDS loading dye (2X) was used and samples were heated at 95 °C for 5 min to elute the bound proteins from the beads. Then eluted fractions were separated on NuPAGE 4-12% Bis-Tris protein gels and transferred to a PVDF membrane through the dry transfer method using the iBlot 2 Dry Blotting System. The membrane was incubated with blocking buffer (5% BSA in TBST) for 1 hour at room temperature before incubation with primary antibodies. The blocked membrane was probed for the following primary antibodies: HA-Tag (C29F4) (Cell Signaling, Cat#3724) (1:2000) and DYKDDDDK Tag (D6W5B) antibody (Cell Signaling, Cat#14793) (1:2000). Primary antibodies were diluted in the same blocking buffer (5% BSA in TBST) and incubated with the membrane at 4 ˚C overnight. The following day, the membrane was washed 3X times with TBST buffer (TBS with 0.1% tween 20) and incubated with the HRP-conjugated secondary antibody (rabbit, Cell Signaling, Cat#7074). Following 1 hour incubation of the secondary antibody at room temperature, the membrane was washed 3 times with TBST buffer, and bands were visualized by chemiluminescence (SuperSignal™ West Pico PLUS Chemiluminescent Substrate, Azure Biosystems C600 imager).

#### 1. 6. Immunoblotting studies to confirm the pyrazole containing PHICS-mediated tyrosine phosphorylation of BRD4 in HEK293T cells.

PHICS-induced tyrosine phosphorylation on BRD4 by ABL1b kinase in HEK293T cells was determined by immunoprecipitation and western blotting. Both kinase and target protein were co-expressed in HEK293T cells before incubation of the compounds. Initially, ABL1b-FLAG and HA-BRD4 plasmids were transfected into HEK293T cells using TransIT®-293 transfection reagent. 36 hours post-transfection, cells were treated with DMSO control and 10 µM of ABL binder, active, and inactive PHICS for 4 hours at 37 °C in DMEM media with the serum. After 4 hours of incubation with compounds cells were washed with cold PBS 1X time and lysed on ice using the cold lysis buffer (M-PER™ Mammalian Protein Extraction Reagent, EDTA-free Protease Inhibitor Cocktail (cOmplete™) and PhosSTOP). Pierce™ BCA Protein Assay Kit was used to determine the protein concentration.

For the immunoprecipitation studies, Anti-HA magnetic beads were used and the immunoprecipitation protocol from Thermo Scientific Pierce was followed. Briefly, 25 µL of beads were taken to the experiment, and beads were equilibrated in lysis buffer before the introduction of the cell lysate. Cell lysates containing one milligram of protein (from each condition) were incubated with the beads at 4 ˚C on a rotating shaker. After overnight incubation, beads were washed 3X times with TBST (0.05% tween 20) buffer and one time with water as per the company protocol before elution of the proteins. SDS loading dye (non-reducing, 2X) was used, and samples were heated at 95 °C for 5 min to elute the bound proteins from the beads. Then eluted fractions were resolved on NuPAGE 4-12% Bis-Tris protein gels and transferred to a PVDF membrane through the dry transfer method using the iBlot 2 Dry Blotting System. After that, the membrane probed for the tyrosine phosphorylation was blocked with 5% milk in TBST, while the membranes probed for the HA and FLAG were blocked with 5% BSA in TBST at room temperature for 1 hour. Then membranes were incubated with the following primary antibodies overnight in a cold room on a rotating shaker: anti-phosphotyrosine antibody, clone 4G10 primary antibody (Millipore Sigma, Cat#05-321) (1:1000), HA-Tag (C29F4) (Cell Signaling, Cat#3724) (1:2000) and DYKDDDDK Tag (D6W5B) antibody (Cell Signaling, Cat# 14793) (1:2000). Next day, membranes were washed with TBST buffer 5 times and incubated with the HRP-conjugated secondary antibodies for 1 hour at room temperature. The membrane probed for the tyrosine phosphorylation was incubated with rabbit anti-mouse IgG (Light Chain Specific) (D3V2A) (Cell Signaling, Cat#58802) (1:1000 dilution). In contrast, the membrane probed for HA and FLAG were incubated with rabbit secondary antibody (Cell Signaling, Cat#7074). After 1 hour of incubation, membranes were washed 3X times with TBST buffer, and bands were visualized by chemiluminescence (SuperSignal™ West Pico PLUS Chemiluminescent Substrate, Azure Biosystems C600 imager).

#### 1. 7. Immunoblotting studies to confirm the DPH-containing PHICS mediate tyrosine phosphorylation of BRD4 in HEK293T cells.

Immunoprecipitation followed by western blotting was performed to detect the proximity-mediated tyrosine phosphorylation on BRD4 by BCR-ABL kinase in HEK293T cells after overexpression of both the kinase and target protein. Initially, BCR-ABL (p210 pcDNA3) and truncated HA-BRD4 plasmids, were transfected into HEK293T cells using TransIT®-293 transfection reagent. 36 hours after of transfection, cells were treated with DMSO control and 10 µM of ABL binder, and DPH-containing PHICS with different exit vectors at 37 °C in DMEM media with the serum. After 4 hours of incubation cells were washed with cold PBS and lysed on ice using the cold lysis buffer (M-PER™ Mammalian Protein Extraction Reagent, EDTA-free Protease Inhibitor Cocktail (cOmplete™) and PhosSTOP). Pierce™ BCA Protein Assay Kit was used to determine the protein concentration.

Immunoprecipitation of HA-tagged BRD4 was performed using anti-HA magnetic beads and the protocol described in section 2.5. Briefly, cell lysates containing one milligram of protein (from each condition) were incubated with 25 µL beads at 4 ºC on a rotating shaker. After overnight incubation, beads were washed three times with TBST buffer and one time with water. SDS loading dye (non-reducing, 2X) was added to washed beads and samples were heated at 95 °C for 5 min to elute the bound proteins from the beads. Then eluted fractions were resolved on Novex Tris-Glycine Gels and transferred to a PVDF membrane using the dry transfer method. Next, the membranes were blocked with 5% milk (for phosphorylation) or 5% BSA (for HA detection) in TBST and incubated overnight in the cold room with the following primary antibodies: anti-phosphotyrosine antibody, clone 4G10 primary antibody (Millipore Sigma, Cat#05-321) (1:1000) and HA-Tag (C29F4) (Cell Signaling, Cat#3724) (1:2000). After washing membranes with TBST buffer five times, membranes were incubated with secondary antibodies from Cell Signaling: Cat#58802 to detect the tyrosine phosphorylation on BRD4 or Cat#7074 to detect the loading levels of HA-BRD4 protein. After 1-hour incubation at room temperature, membranes were washed three times with TBST buffer, and bands were visualized by chemiluminescence (SuperSignal™ West Pico PLUS Chemiluminescent Substrate, Azure Biosystems C600 imager).

### 1. 8. Immunoblotting studies to confirm that pyrazole containing PHICS mediated tyrosine phosphorylation of EGFR and HER2 expressed in HEK293T cells.

HEK293T cells were seeded into a 24-well plate at the density of 0.25 million cells per well one day prior to the transfection of ABL-HA with either inactive EGFR (iEGFR-FKBP-FLAG) or with active HER2 (HER2-FKBP-FLAG), respectively. 24 hours post-transfection, cells were treated with ABL-FKBP-PHICS at 500nM concentration for 1 hour, washed with PBS, harvested, and lysed using lysis buffer (MPER buffer (cat#78505, Thermo Scientific) containing protease (cat#4693159001, Roche) and phosphatase inhibitor (ca#4906837001, Roche) cocktails). The total lysate was spun at 16 000 g for 10 min at 4 °C to separate the supernatant. Total protein concentration was estimated by BCA**.** Cell lysates containing equal amounts of the total protein (25-30 µg) were heated with Laemmli SDS sample buffer (reducing) at 95-100 °C for 5min. The protein samples were then resolved by NuPAGE gels and transferred to nitrocellulose membrane using the dry-transfer method (iBlot2). The membranes were blocked with 5% milk in TBST buffer followed by incubation with primary antibodies (EGFR phospho-antibodies: pY845, cat# 6963, Cell Signaling; pY992, cat#2235, Cell Signaling; pY998, cat#2641, Cell Signaling; pY1045, cat#2237, Cell Signaling; pY1068, cat#2236, Cell Signaling; pY1086, cat#2220, Cell Signaling; pY1148, cat#4404, Cell Signaling; pY1173, cat#4407, Cell Signaling; anti-pTyr-pan, cat# 8945, Cell Signaling; anti-FLAG (mouse), cat# 8146, Cell Signaling; anti-FLAG (rabbit), cat# 14793, Cell Signaling; HER2 phospho-antibodies: pY877, cat#2241, Cell Signaling; pY1196, cat#6942, Cell Signaling; pY1221, cat#2243, Cell Signaling; pY1248, cat#2247, Cell Signaling) overnight at 4 ˚C on a shaker (cold room). Unbound primary antibodies were washed out with TBST buffer and membranes were incubated further with secondary antibodies (IRDye800CW, Donkey anti-Mouse, cat#926-32212; IRDye680RD, Donkey anti-Rabbit, cat#926-68073) for 1 hour at room temperature. Unbound secondary antibodies were washed out with TBST buffer and blots were developed on LiCor Odyssey CLx Imaging System. All primary and secondary antibodies were used at 1:1000 and 1:10,000 dilutions, respectively.

### 1. 9. Immunoblotting studies to confirm PHICS-mediated tyrosine phosphorylation of BRD4 by BTK in HEK293T cells.

HEK293T cells were seeded into a 6-well plate at the density of 1.2 million cells per well one day prior to BTK-FLAG and HA-BDR4 co-transfection. 24 hours post-transfection, cells were treated with BTK-PHICS at 500nM concentration for 4 hours, washed with cold PBS, harvested, and lysed using lysis buffer (MPER buffer (cat#78505, Thermo Scientific) containing protease (cat#4693159001, Roche) and phosphatase inhibitor (ca#4906837001, Roche) cocktails). The total lysate was spun at 16 000 g for 10 min at 4 °C to separate the supernatant. The total protein concentration was estimated by BCA**. HA-IP:** HA-magnetic beads (cat#88837, Thermo Scientific) (10 µL slurry per sample) were taken and equilibrated twice with MPER buffer. An equal amount of the above-prepared supernatant (total protein) was added to the equilibrated beads and incubated with gentle rotation overnight at 4 ˚C (cold room). Unbound protein was washed out using TBST buffer three times, and the bound protein was eluted with 4x SDS sample buffer (20 µL per tube) by heating at 95-100 °C for

5 min. The eluted protein was taken out by magnetic separation to a fresh tube. Equal volumes of pulldown samples were resolved by 4-12% Bis-Tris NuPAGE gels and transferred to nitrocellulose membrane using the dry-transfer method (iBlot2). The membranes were blocked with 5% BSA in TBST followed by incubation with primary antibodies (anti-pTyr-pan, cat#8954, Cell Signaling; anti-HA, cat#2367, Cell Signaling) overnight at 4 ˚C on a shaker (cold room). Unbound primary antibodies were washed out with TBST buffer and membranes were further incubated with secondary antibodies (IRDye800CW, Donkey anti-Mouse, cat#926-32212; IRDye680RD, Donkey anti-Rabbit, cat#926-68073) for 1 hour at room temperature. Unbound secondary antibodies were washed out with TBST buffer and blots were developed on LiCor Odyssey CLx Imaging System. All primary and secondary antibodies were used at 1:1000 and 1:10,000 dilutions, respectively.

#### 1. 10. Immunoblotting studies to confirm PHICS-mediated phosphorylation of Halo-Tag by ABL in HEK293T cells.

HEK293T cells stably expressing Halo-Tag were seeded into a 6-well plate at the density of 1.2 million cells per well one day prior to ABL-HA transfection. 24 hours post-transfection, cells were treated with DMSO, control compounds, or 5 μM of ABL-Halo-Tag PHICS. After 4 hours incubation with compounds, media was removed, cells washed with cold PBS, harvested, and lysed using lysis buffer (MPER buffer (cat#78505, Thermo Scientific) containing protease (cat#4693159001, Roche) and phosphatase inhibitor (ca#4906837001, Roche) cocktails). The total lysate was spun at 16 000 g for 10 min at 4 °C to separate the supernatant. The total protein concentration was estimated by BCA**. FLAG-IP:** FLAG-magnetic beads (cat#A36798, Thermo Scientific) (10 μL slurry per sample) were taken and equilibrated twice with MPER buffer. Equal amounts of the above-prepared supernatants (total protein) were added to the equilibrated beads and incubated with gentle rotation overnight at 4 ˚C (cold room). Unbound protein was washed out using TBST buffer three times, and the bound protein was eluted with 4x SDS sample buffer (20 μL per tube) by heating at 95-100 °C for 5 min. The eluted protein was taken out by magnetic separation to a fresh tube. Equal volumes of pulldown samples were resolved by 4-12% Bis-Tris NuPAGE gels and transferred to nitrocellulose membrane using the dry-transfer method (iBlot2). The membranes were blocked with 5% BSA in TBST followed by incubation with primary antibodies (anti-pTyr-pan, cat#8954, Cell Signaling; anti-FLAG (mouse), cat#8146, Cell Signaling) overnight at 4 ˚C on a shaker (cold room). Unbound primary antibodies were washed out with TBST buffer and membranes were further incubated with secondary antibodies (IRDye800CW, Donkey anti-Mouse, cat#926-32212; IRDye680RD, Donkey anti-Rabbit, cat#926-68073) for 1 hour at room temperature. Unbound secondary antibodies were washed out with TBST buffer and blots were developed on LiCor Odyssey CLx Imaging System. All primary and secondary antibodies were used at 1:1000 and 1:10,000 dilutions, respectively.

#### 1. 11. Immunoblotting studies to confirm PHICS-mediated tyrosine phosphorylation in HEK293T cells with transiently overexpressed BTK-FLAG and ABL-HA.

HEK293T cells were seeded into a 6-well plate at the density of 2 million cells per well in 2 mL of media one day prior to BTK-FLAG and ABL-HA co-transfection (1 μg of each DNA in 200 μL OPTI-MEM with 6 μL of trans-IT per well). 24 hours post-transfection, cells were treated with BTK-ABL PHICS or a mixture of binders at 1 μM concentration; 2 wells were used per one condition. After 4 hours of incubation, the media was removed. Cells were washed with 2 mL of ice-cold PBS and lysed using 350 µL of lysis buffer per well (MPER buffer (cat#78505, Thermo Scientific,) containing protease (cat#4693159001, Roche, 1 tablet per 7 mL of buffer) and phosphatase inhibitor (ca#4906837001, Roche, 1 tablet per 7 mL of buffer) cocktails). The total lysate was spun at 16 000 g for 10 min at 4 °C to separate the supernatant. Total protein concentration was estimated by BCA, and concentrations were adjusted to be equal by the addition of lysis buffer to samples with higher concentrations. **HA-IP:** HA-magnetic beads (cat#88837, Thermo Scientific) (25 µL slurry per sample) were taken and equilibrated twice with TBST buffer (1 mL). The equal volumes of supernatant containing 1.2 mg of protein were added to the equilibrated beads and incubated with gentle rotation overnight at 4 ˚C (cold room). Unbound protein was washed out using TBST buffer (300 µL) three times and distilled water (300 µL) two times. The bound protein was eluted with reducing 2x Laemmli SDS sample buffer (AlfaAesar, J61337) (40 µL per tube) by heating at 95-100 ºC for 5 min. The eluted protein was taken out by magnetic separation to a fresh tube. Equal volumes (5 µL) of pulldown samples were resolved by 4-12% Bis-Tris NuPAGE gels and transferred to the PVDF membrane using the dry-transfer method (iBlot2). The membranes were blocked with 5% milk in TBST buffer, followed by incubation with primary antibodies (anti-pTyr-pan, cat#8954, Cell Signaling; anti-HA, cat#2367, Cell Signaling) overnight at 4 ˚C on a shaker (cold room). Unbound primary antibodies were washed out with TBST buffer and membranes were further incubated with secondary antibodies (IRDye800CW, Donkey anti-Mouse, cat#926-32212; IRDye680RD, Donkey anti-Rabbit, cat#926-68073) for 1 hour at room temperature. Unbound secondary antibodies were washed out with TBST buffer five times, and blots were developed using the LI-COR system. **FLAG-IP:** FLAG-magnetic beads (cat#A36798, Thermo Scientific) (50µL slurry per sample) were taken and washed with MPER buffer (500 µL) three times. Equal volumes of supernatant containing 1.2 mg of protein were added to the washed beads and incubated with gentle rotation overnight at $4^0$C (cold room). Unbound protein/flow-through was washed out using 1xPBS buffer twice (500 µL) and distilled water (1 mL) twice. The bound protein was eluted with reducing 2x Laemmli SDS sample buffer (AlfaAesar, J61337) (40µL per tube) by heating at 95-100 ºC for 5min. The eluted protein was taken out by magnetic separation to a fresh tube. The pulldown samples' equal volumes (5 µL) were resolved by nuPAGE gels and transferred to the PVDF membrane using the dry-transfer method (iBlot2). The membranes were blocked with 5% milk in TBST buffer, followed by incubation with primary antibodies (anti-pTyr-pan, cat#8954, Cell Signaling; anti-FLAG, cat#F3165, Sigma Aldrich) overnight at $4^0$C on a shaker (cold room). Unbound primary antibodies were washed out with 1xTBST buffer and incubated further with secondary antibodies (IRDye800CW, Donkey anti-Mouse, cat#926-32212; IRDye680RD, Donkey anti-Rabbit, cat#926-68073) for 1 hour at room temperature. Unbound secondary antibodies were washed out with TBST buffer and blots were developed on LiCor Odyssey CLx Imaging System. All primary and secondary antibodies were used at 1:1000 and 1:10,000 dilutions, respectively.

#### 1. 12. Cell viability studies with PHICS.

Cell viability studies were performed in 384 well plates (Corning 3826). Compounds and DMSO control were printed into wells (avoiding edge wells) using Tecan D300e Digital Dispenser with each concentration being performed in triplicate. 2000 cells (Mino cells, ATCC CRL-3000; Raji cells, ATCC CCL-86; K562s cells, ATCC CRL-3343) were added in 50 µL of media to each well with compounds. Several wells were filled with media for the estimation of the background signal. Plates with cells and compounds were incubated for 72 hours and treated with Promega's Cell Titer Glo reagent (40 µL per well) following the procedure provided by the vendor. After 30 min incubation at room temperature, the luminescence signal was recorded using a Perkin Elmer Envision plate reader. Data were analyzed by subtracting the average signal in media-containing wells from the signal in compound-treated wells. Viability in each compound-treated well was calculated by dividing the signal in compound-treated wells by the average signal in DMSO-treated wells and multiplying by 100%. Dose curves and $EC_{50}$ values were obtained using Graph PRISM9 software.

### 1. 13. Labeling of BTK in cells and its effect on autophosphorylation of BTK.

#### 1. 13. 1. Cell treatment.

HEK293T cells were seeded into a 6-well plate at the density of 2 million cells per well in 2 mL of media one day prior to BTK-FLAG transfection (2 µg of DNA in 200 µL OPTI-MEM with 6 µL of trans-IT per well). 24 hours post-transfection, cells were treated with methacrylamides **13-16**, BTK-ABL PHICS **17** and **18**, and control compounds (DMSO, amines **40** and **45**) at 1 µM concentration. After 4 hours of incubation with compounds, the media was removed. Cells were washed with 2 mL of ice-cold PBS and lysed using 350 µL of lysis buffer per well (MPER buffer (cat#78505, Thermo Scientific,) containing protease (cat#4693159001, Roche, 1 tablet per 7 mL of buffer) and phosphatase inhibitor (ca#4906837001, Roche, 1 tablet per 7 mL of buffer) cocktails). The total lysate was sonicated for 15 seconds and spun at 16 000 g for 10 min at 4 ºC to separate the supernatant. Total protein concentration was estimated by BCA, and concentrations were adjusted to be equal by adding lysis buffer to samples with higher concentrations**. In-gel fluorescence:** To demonstrate the labeling of BTK with alkyne-containing methacrylamides, copper-catalyzed click reaction with sulfoCy5.5 azide and in-gel fluorescence was performed. Click reaction was performed in the total volume of 30 µL by the addition of 1.5 µL of 800 µM solution of sulfoCy5.5 azide (BroadPharm, cat# BP-22483), 1.5 µL of 60 mM solution of tris(3-hydroxypropyltriazolylmethyl)amine (THPTA, Lumiprobe, cat# H4050), 1.5 µL of 60 mM $CuSO_4$, and 1.5 µL of 60 mM solution of 74 mM sodium ascorbate to 24 µL of lysates obtained after treatment with DMSO, **15**, and **16.** After 30 min incubation at room temperature, 10 µL of 4x Laemmli SDS-sample reducing buffer (bioWorld, cat# 10570020) was added, and samples were heated at 100 ºC for 5 min. After cooling to room temperature, 20 µL of the reaction mixtures were resolved on NuPAGE 4-12% Bis-Tris gels (ThermoFisher, cat # NP0321BOX) with MOPS SDS running buffer (ThermoFisher, cat # NP0001) at 200V over 1 hour. The gels were visualized using Azur c600 Gel Imaging system by Biosystems at 700 nm (NIR).

#### 1. 13. 2. Effect of compounds on autophosphorylation of BTK.

10 µL of lysates obtained after treatment with tested compounds were incubated with 3.5 µL of 4x Laemmli SDS-sample reducing buffer (bioWorld, cat # 10570020) at 100 ºC for 5 min. After cooling to room temperature, samples were resolved on NuPAGE 4-12% Bis-Tris gels (ThermoFisher, cat # NP0321BOX) with MOPS SDS running buffer (ThermoFisher, cat # NP0001) at 200V over 1 hour and transferred to PVDF membrane (ThermoFisher, cat # IB24002) using the dry-transfer method (iBlot2). The membranes were blocked with 5%

BSA in TBST buffer over 1 hour at room temperature, followed by incubation with primary antibodies (anti-FLAG antibody from Sigma (cat #F1804) and phospho-BTK (Tyr223) antibody from Cell Signaling (cat #5082)) overnight at 40 ºC on a shaker. Unbound primary antibodies were washed out with TBST buffer five times and incubated further with secondary antibodies (IRDye800CW, Goat anti-Rabbit, cat#926-32211; IRDye680RD, Goat anti-Mouse, cat#926-68070 from LI-COR) for 1 hour at room temperature. Unbound secondary antibodies were washed out with TBST buffer five times, and the resulting blot was developed on LiCor Odyssey CLx Imaging System.

**1. 14. SRE reporter assay for EGFR signaling.**

HEK293 cells were seeded in a 6-well plate at a density of 1.2 million cells per well in 2 ml of DMEM media with 10 % FBS. The next day, the cells were transfected with 100 ng EGFR-BTB-FKBP$^{F36V}$ and 1 μg SRE plasmids using TransIT-LT1 (Mirus) transfection reagent. After 6 hrs of transfection, the cells were harvested and plated at a density of 40,000 cells per well in a white 96-well plate in 100 μl opti-MEM media per well. The cells were starved overnight in opti-MEM media to remove any signaling events resulting from the serum. Further, the cells were treated with 50 nM of monofunctional (Abl1 binder) and bifunctional (Abl1 - FKBP$^{F36V}$) molecules along with 1 μM of BI-3802 (BTB$_{BCL6}$ dimerizer), to induce dimerization of EGFR keeping DMSO as a control. The cells were incubated for 7 hrs after treatment. The phosphorylation of EGFR by endogenous Abl proteins results in downstream signaling, activation, and expression of the reporter luciferase gene placed under the SRE promoter. The increase in luminescence was measured following the protocol mentioned for the ONE-Glo™ luciferase assay system (Promega) using PerkinElmer EnVision multimode plate reader.

**1. 15. NanoBRET Intracellular kinase target engagement assay.**

NanoBRET assay was performed according to the manufacturer's protocol (Promega cat#N2641). HEK293T cells were washed with PBS, trypsinized, and centrifuged for 5 minutes at 200 g. Transfection of carrier DNA (Promega cat#N2641) and BTK Kinase-NanocLuc fusion vector (Promega cat#N2441) were both diluted to 9 µg/ml and 1 µg/ml respectively, in 1 ml of Opti-MEM I (Life Technologies cat# 11058-021) no phenol red, 1 % FBS) accordingly. 30µl of FuGENE HD Transfection Reagent (Promega cat# E2311)) was added to 1 ml of DNA mixtures, the solution was mixed by inversion 5-10 times and incubated at room temperature for 20 minutes, after which the mix was added to 20 ml of HEK293T cell suspension of 2*10$^5$ cell/ml in Opti-MEM assay solution. The resulting solution was gently mixed by inversion five times, and 100 µl of the mixture was dispensed in each well of the 96-well plate (20000 cells/well, Corning cat# 3917). After 24 h, compound dilutions of 10 µl at 10x of tested concentrations were added to corresponding wells in triplicates, and 5 µl of 20x of NanoBRET K-10 tracer (Promega cat#N2641) solution prepared by mixing 1 part of 100x K-10 tracer with 4 parts of Tracer Dilution Buffer. The final K-10 tracer concentration in each well was 0.5 µM. After 2 h incubation at 37 °C in a 5% $CO_2$ humidified atmosphere, the plate was equilibrated at room temp for 15 minutes. After equilibration, 50 µl of 3X Complete Substrate plus Inhibitor Solution (3 equiv. of NanoBRET™ Nano-Glo. Substrate and 1 equiv. of Extracellular NanoLuc. Inhibitor from the assay kit) was added to each well and incubated for 2 minutes. Both Donor emission wavelength (450 nm) and acceptor emission (610 nm) were measured by PerkinElmer EnVision multimode plate reader following Promega's protocol suggestions. To obtain final BRET ratios (mBU), the acceptor emission value was divided by the donor emission value for each sample and multiplied by 1000.

### 2. Supplemental figures.

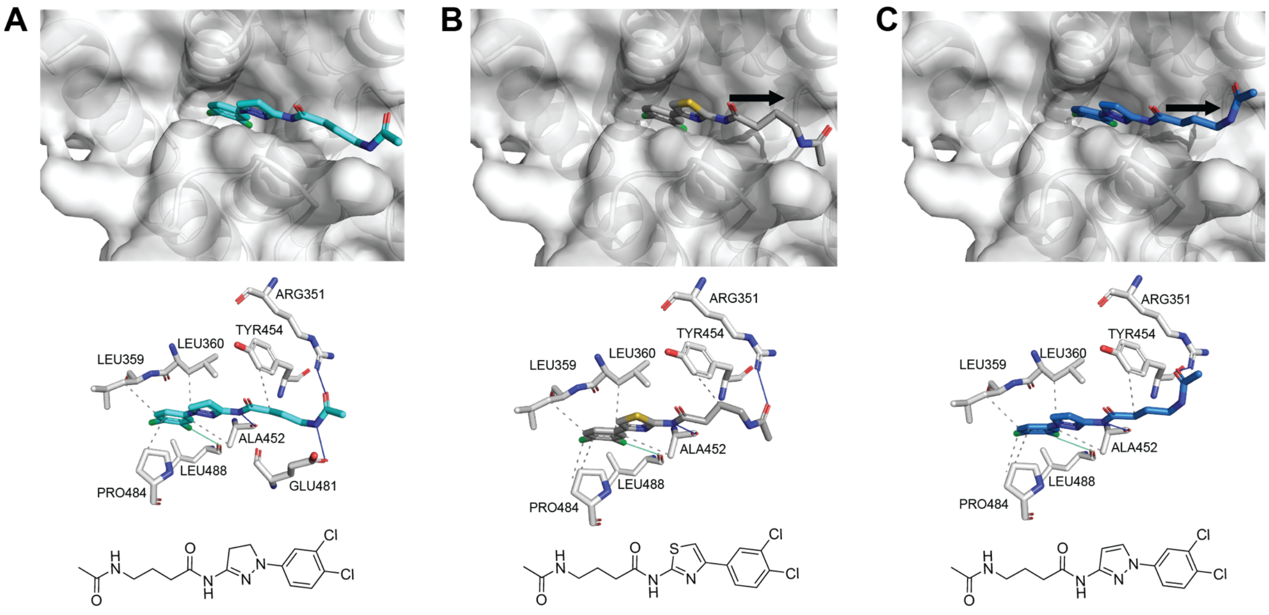

**Figure S1. (A-C)** Docking studies with the first class of ABL binders. Identification of linker attachment sites on dihydropyrazole **(A)**, thiazole **(B)** and pyrazole **(C)** ABL binders.

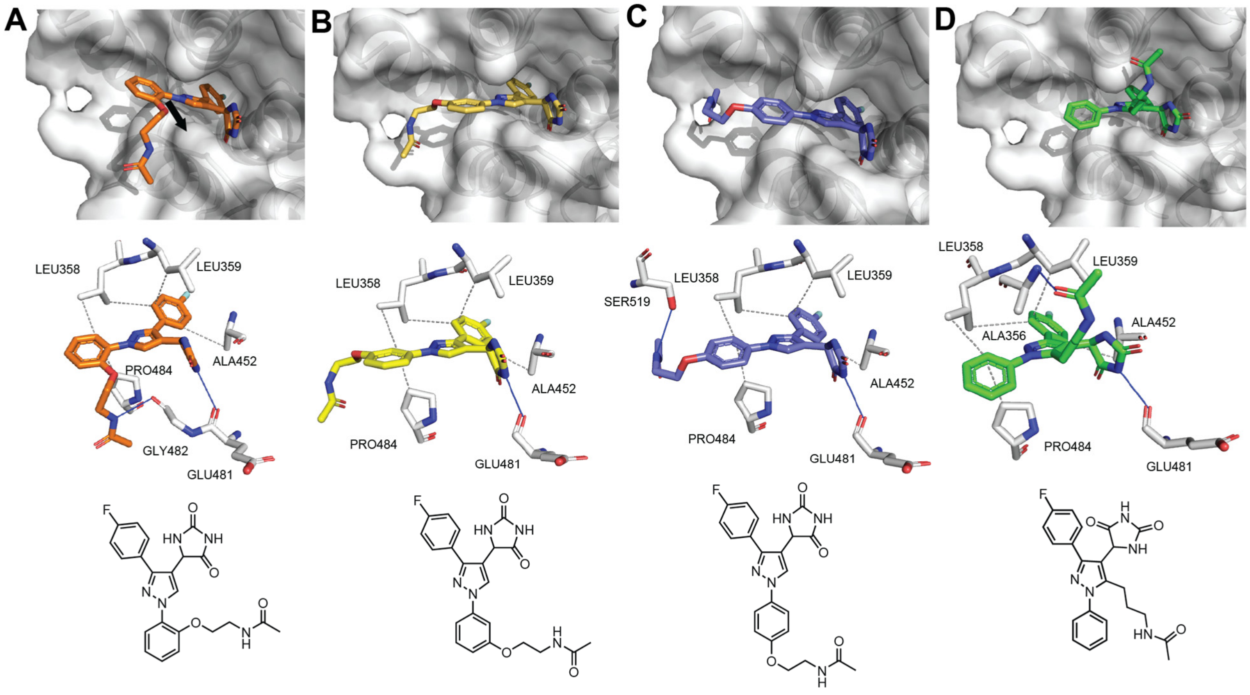

**Figure S2. (A-D)** Docking studies with the second class of ABL binders. Identification of linker attachment sites on DPH core: ortho- **(A),** meta- **(B),** para- **(C),** and pyrazole **(D)** exit vectors.

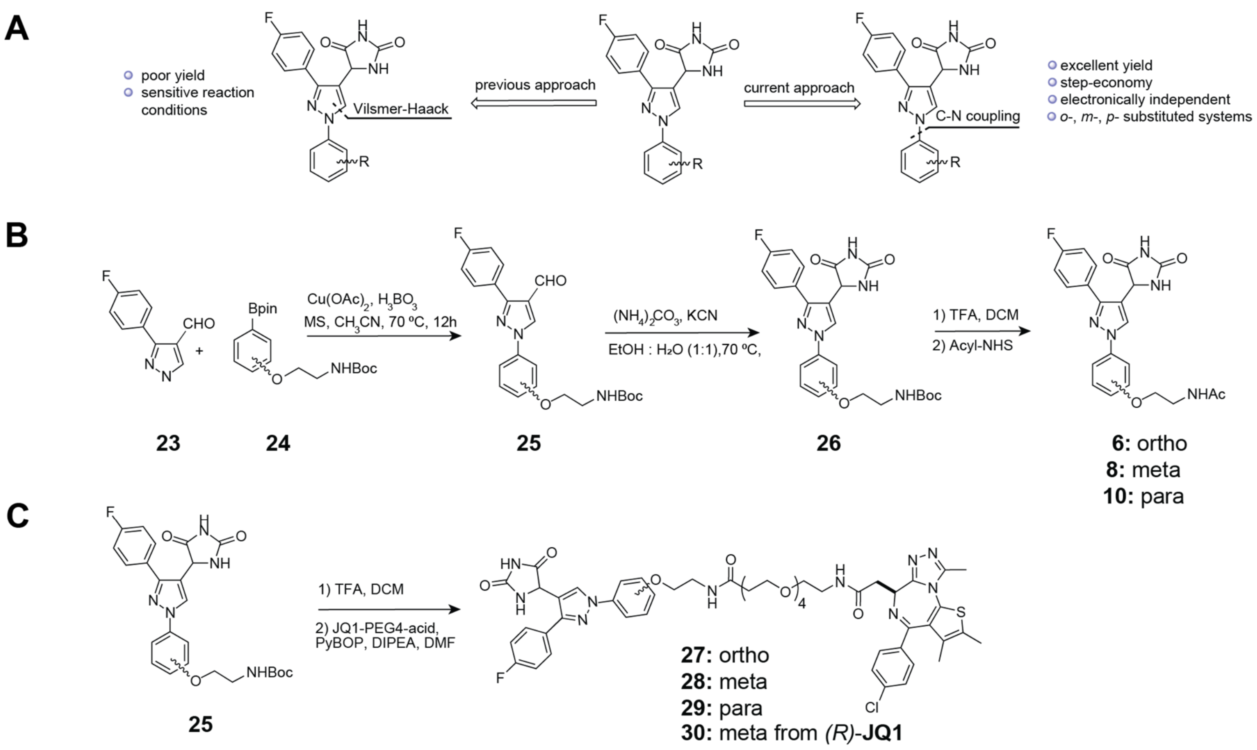

**Figure S3. (A)** Comparison of current and previous approaches of ABL binder synthesis based on hydantoin. **(B)** Synthesis of hydantoin-derived ABL binders with ortho-, meta- and para- exit vectors. **(C)** Synthesis of BRD4-targeting ABL PHICS based on hydantoin binder with ortho-, meta- and para- exit vectors.

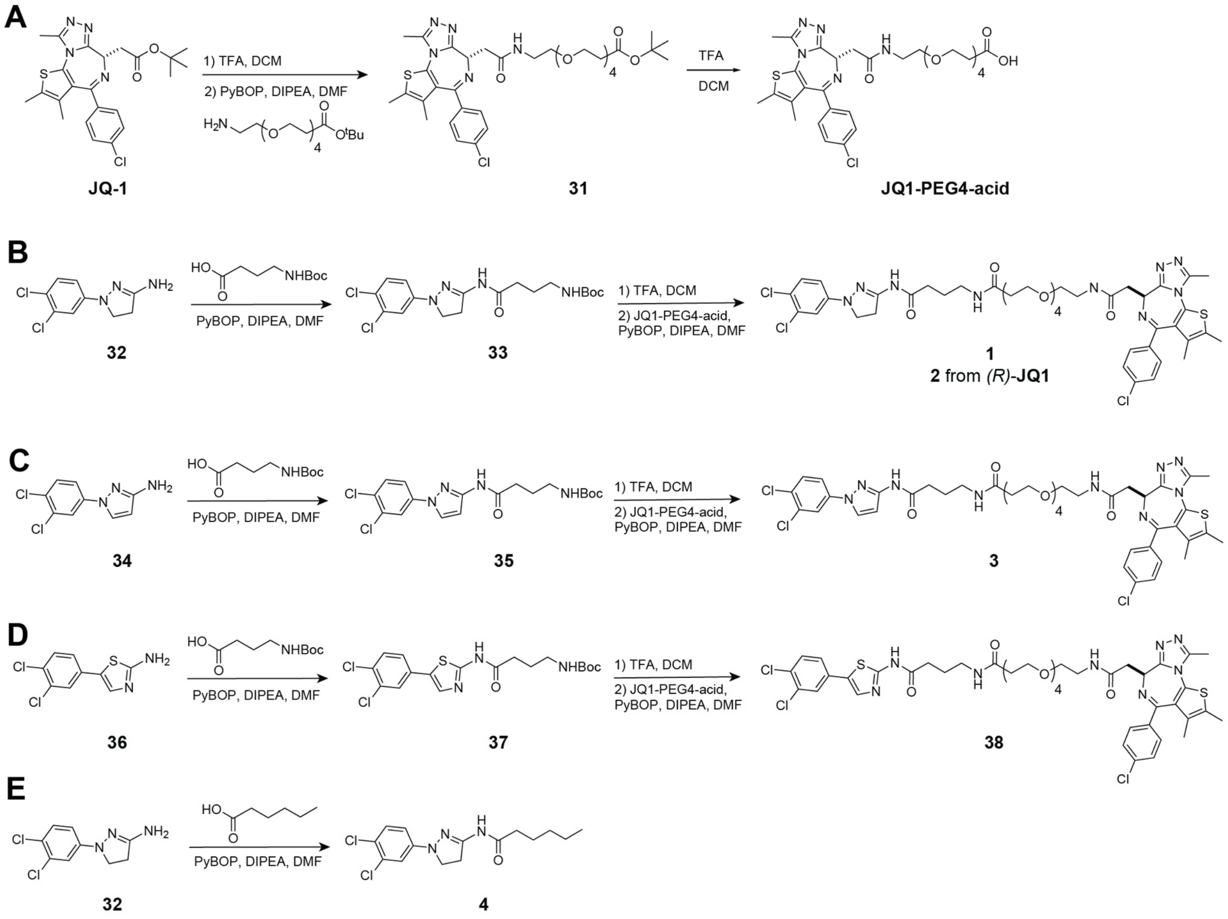

**Figure S4. (A)** Synthesis of JQ1-PEG4-acid. **(B-D)** Synthesis of BRD4-targeting ABL PHICS based on dihydropyrazole **(B)**, pyrazole **(C-)**, and thiazole **(D)** binders. **(E)** Synthesis of ABL binder control.

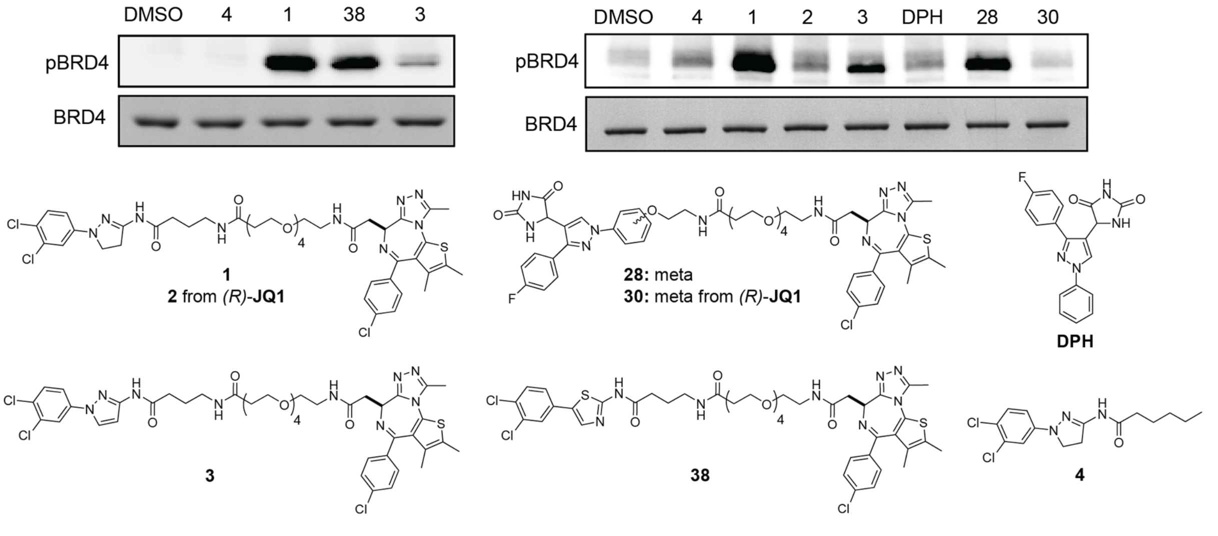

**Figure S5.** Identification of the optimal PHICS for induction of BRD4 phosphorylation by ABL in vitro.

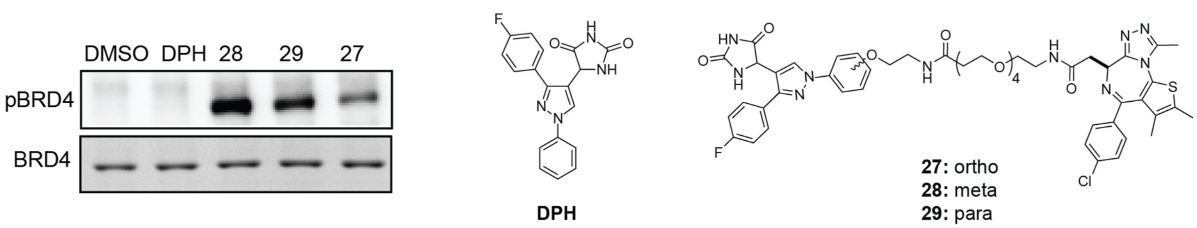

**Figure S6.** Identification of the optimal exit vector for DPH-derived BRD4-ABL PHICS.

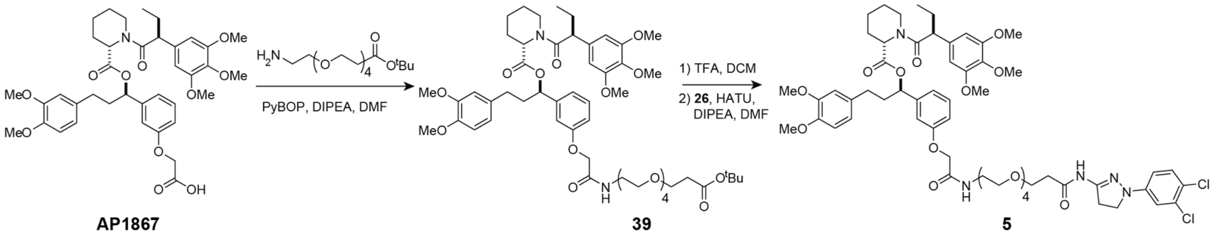

**Figure S7.** Synthesis of FKBP$^{F36V}$ -ABL PHICS.

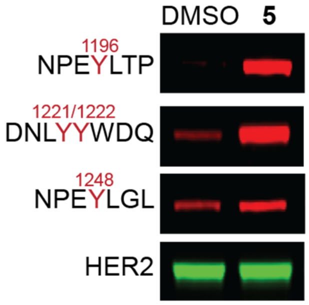

**Figure S8.** Phosphorylation of catalytically inactive HER2-FKBP$^{F36V}$ by ABL in cells in the presence of **5** detected using phospho-HER2 specific antibodies.

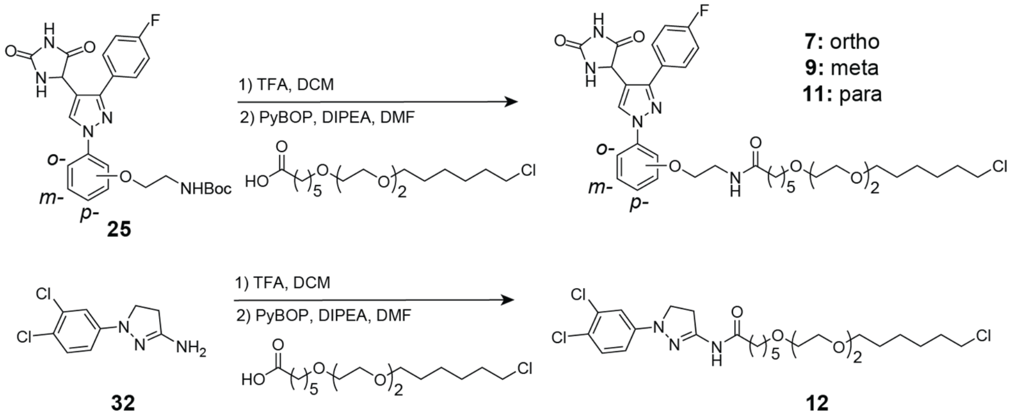

**Figure S9.** Synthesis of ABL-HaloTag PHICS.

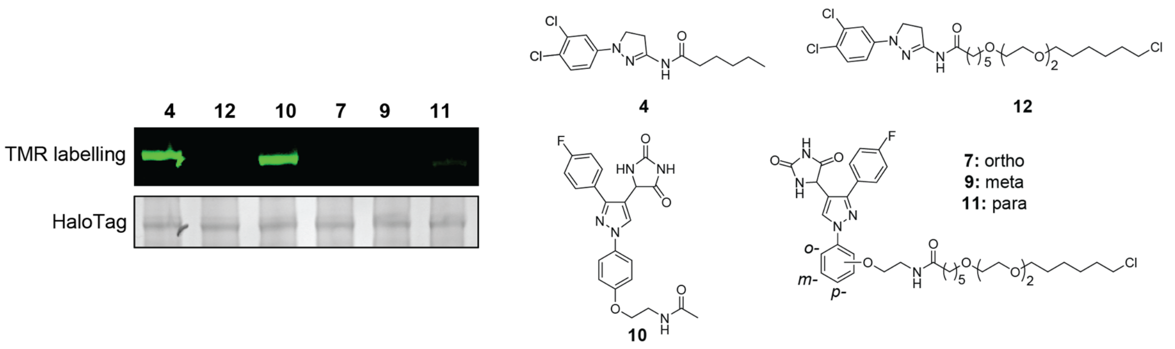

**Figure S10.** Evaluation of cell-permeability of ABL-HaloTag PHICS and efficiency of HaloTag labeling with TMR- HaloTag probe.

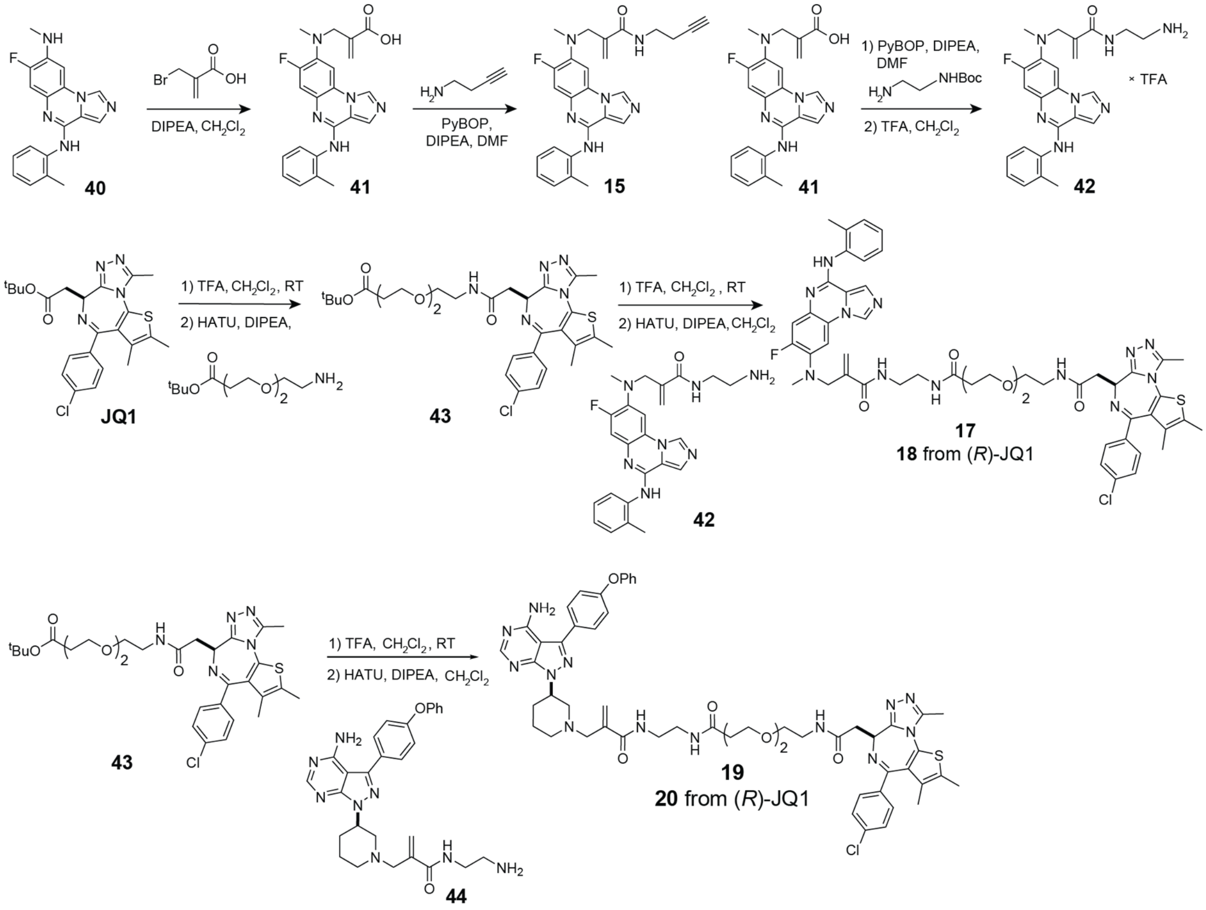

**Figure S11.** Synthesis of **8** and BTK-BRD4 PHICS.

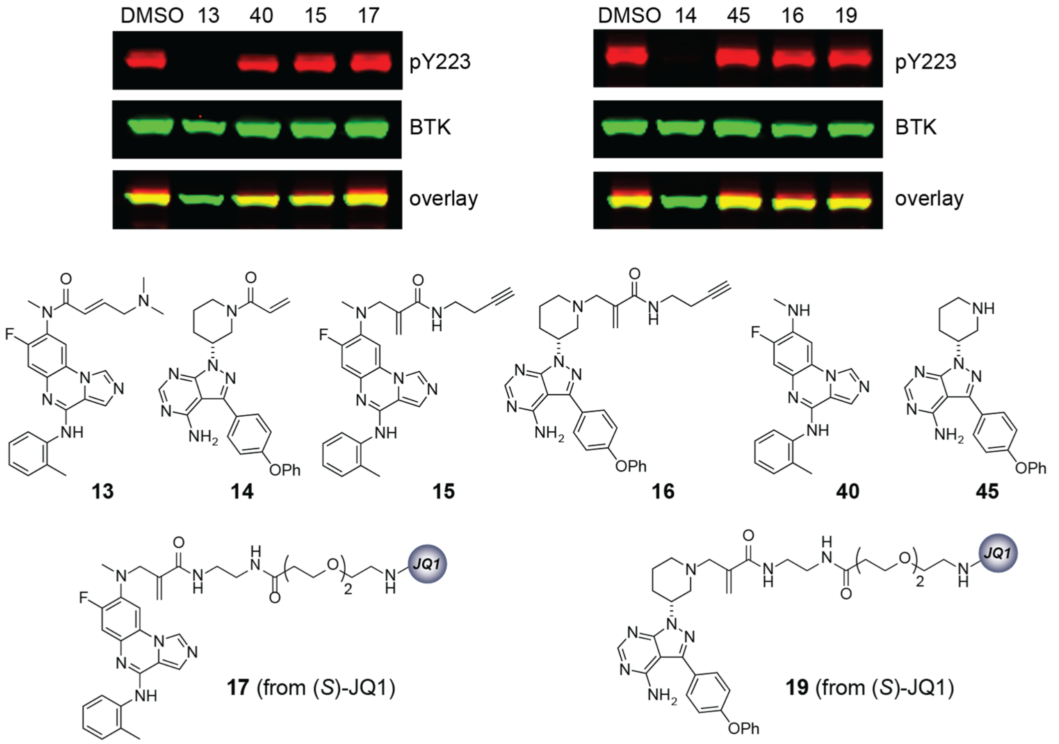

**Figure S12.** Effect of acrylamide inhibitors **13** and **14**, alkyne-containing probes **15** and **16**, and BTK-BRD4 PHICS **17** and **19** on autophosphorylation level of BTK.

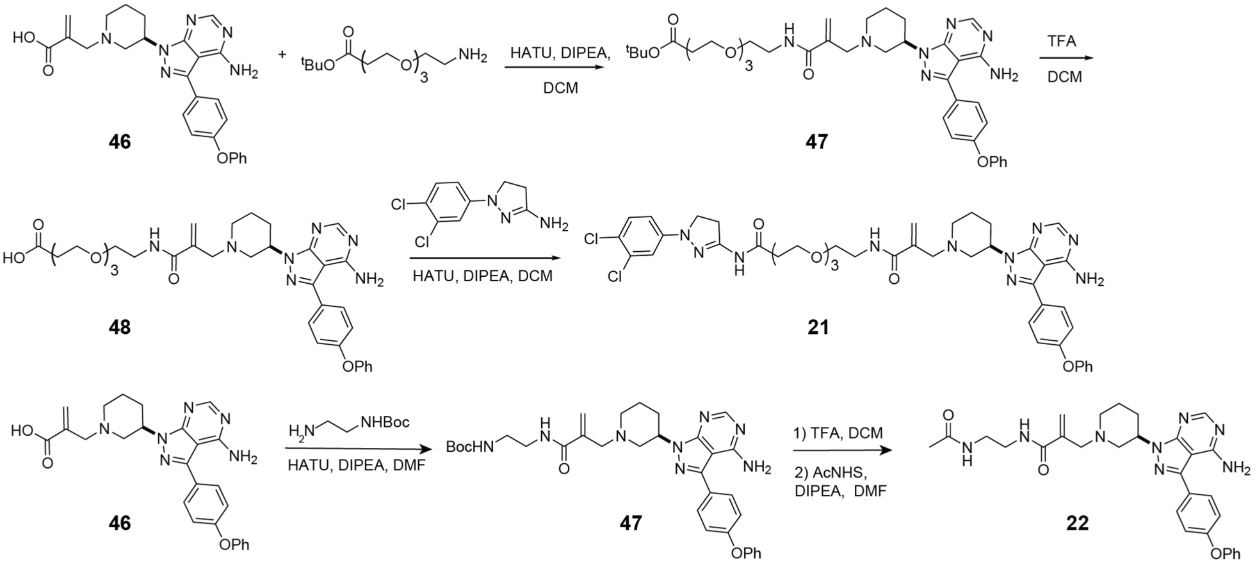

**Figure S13.** Synthesis of ABL-BTK PHICS and BTK-binder control.

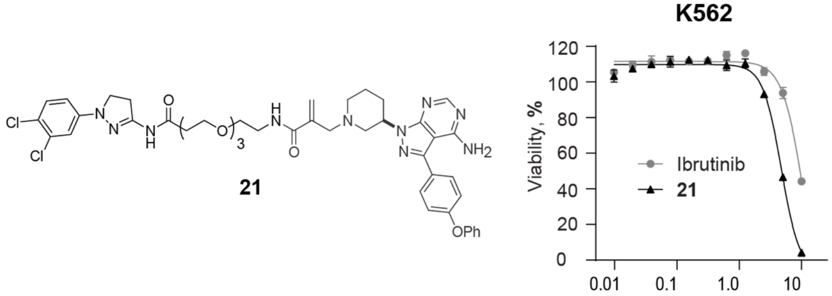

**Figure S14.** Effect of PHICS **21** on the viability of K562 cells.
